# Low frequency oscillatory activity of the subthalamic nucleus is a predictive biomarker of compulsive-like cocaine seeking

**DOI:** 10.1101/451203

**Authors:** Mickaël Degoulet, Alix Tiran-Cappello, Christelle Baunez, Yann Pelloux

## Abstract

Cocaine seeking despite a foot-shock contingency is used to model compulsive drug seeking, a core component of drug addiction, in rodents. Deep brain stimulation (DBS) of the subthalamic nucleus (STN) is efficient on other addiction criteria models and we show here that 30-Hz STN stimulation reduces pathological cocaine seeking in compulsive-like rats. This confirms STN DBS as a potential strategy to treat addiction. We also observed that only ‘compulsive-like’ rats displayed a progressive increase in STN low frequency oscillations, especially in the alpha/theta band (6-13 Hz), during cocaine escalation. Conversely, applying 8-Hz STN DBS to mimic alpha/theta oscillations in ‘non-compulsive’ animals changed them into ‘compulsive’ ones. We have thus identified a predictive neuronal biomarker of compulsivity. Since one critical challenge in addiction research is to identify vulnerable individuals before they transition to harmful drug consumption pattern, our results could lead to new diagnostic tools and prevention strategies.

## Introduction

Among drug users, only a small fraction of individuals transitions to addiction through loss of control over their drug intake and continues to seek drugs of abuse despite obvious negative consequences for their personal, professional and social life (*1*). Yet, identifying predictive neuronal biomarkers in these individuals remains a key challenge for both the prevention and the treatment of addictive disorders. In a rat model of compulsivity, perseverance of drug seeking-taking behaviors despite negative consequences can be observed by associating a punishment (*i.e.* mild electric foot shock) to seeking/consumption paradigms, thereby offering a relevant model to study one of the core features of drug addiction in humans (*2*, *3*)). Indeed, prior studies have shown that, in rats that had long term exposure to cocaine self-administration, a subset of animals displays compulsive-like cocaine seeking and taking behaviors despite the foot-shock punishment (*4*–*9*). This allowed critical advances in characterizing brain cellular and molecular alterations consequent to extended cocaine access and exposure to punishment paradigms, notably neuronal activity and neuroplasticity (*4*, *10*–*13*). However, addiction research has failed so far to identify alterations in brain region activity preceding behavioral expression of compulsive-like seeking/taking behaviors. Such predictive biomarkers would be of critical importance to help identifying subjects susceptible to develop more harmful patterns of consumption.

The subthalamic nucleus (STN), which is the current target for surgical treatment of Parkinson’s disease (PD), displays neurophysiological signatures, especially in terms of abnormal neural oscillations, in PD (*14*, *15*)) and obsessive-compulsive disorder (OCD) patients, with a specific oscillatory activity related to compulsive behaviors (*16*, *17*)). Notably, PD patients suffering from impulse control disorders exhibit an increased alpha-theta (6-13 Hz) activity in the STN when receiving dopaminergic agonist treatment (*18*). These data suggest that the STN could be a potential place to look for a predictive marker of compulsive drug use. In the context of addiction indeed, the STN has recently received much attention since its lesions or manipulation of its activity with high frequency (130-Hz) deep brain stimulation (DBS) not only decreases the rat’s motivation to work for cocaine, heroin or alcohol (*19*–*21*) but also prevents escalation of cocaine intake and further reduces cocaine, heroin and alcohol re-escalation normally observed after a period of protracted abstinence (*20*, *22*, *23*). Particularly relevant to the present study, the development of cocaine escalation induces a progressive increase in STN low frequency oscillations, measured with local field potentials (LFPs), compared to its activity during short access to cocaine (*22*). This suggests a key contribution of the STN oscillatory activity to the development of loss of control over drug intake, which allows the emergence of compulsive-like seeking behaviors (*5*, *9*)). We thus hypothesized here that STN oscillatory activity might be involved in the onset of compulsive drug seeking and thus performed LFPs recordings, through DBS electrodes implanted in the STN during the cocaine escalation protocol, before subjecting the animals to the punished seeking paradigm.

## Results

### Compulsive-like cocaine seeking emerges following escalation of cocaine intake

To investigate the involvement of STN activity in compulsive-like seeking behaviors, we used a sequential model in which a subset of animals with an extended cocaine self-administration history keeps seeking the drug despite intermittent punishment by an electric foot shock. Briefly (see methods for details), animals were trained to press a seeking lever at random intervals to access the second stage of the seeking cycle, which triggers retraction of the seeking lever and insertion of the cocaine-taking lever. One press on this lever triggers *i.v.* cocaine delivery followed by a 10-min time-out period. At the end of the training, animals could complete up to 10-daily seeking cycles (*i.e.* 10 cocaine infusions).

After this conditioning, the control animals (n = 9) were immediately subjected to eight punished sessions. Here, 50% of the seeking cycles ended with the pseudorandom delivery of a mild electric foot shock (0.5 mA, 0.5 sec) with no access to the taking lever, in contrast to the other half that allow cocaine injections. In this condition, all animals quickly ceased seeking cocaine when exposed to the punishment contingency (Fig. S1A: *F*_*(12,96)*_ = 90.66, *P* < 0.0001).

The other animals (N = 53) were subjected to 6 h-daily access to cocaine under a fixed ratio 1 schedule of reinforcement (FR1) for 15 days, leading to an escalation in drug intake (*24*) (Fig. S1B: session effect: *F*_(*14,714)*_ = 6.71, *P* < 0.0001), before being exposed to the punishment contingency (Fig. 1A). After escalation of drug intake, the animals were tested again in the seeking/taking task for five baseline seeking sessions before the eight punished sessions. Previous reports (*4*, *5*, *7*, *8*) indicate that the population distribution of cocaine seeking/taking behaviors under punishment is bimodal: one ‘compact’ population of low responders (*i.e.* ‘shock-sensitive’, ~2/3 of the population) and a second one more ‘broadly-distributed’ that keeps seeking drug despite punishment (*i.e.* ‘shock-resistant’, ~1/3 of the population). Frequency histogram analysis of the sum of seeking cycle completed during the last four punishment sessions clearly identifies ‘shock-sensitive’ (n =36) and ‘shock-resistant’ (n = 17) animals (Fig. S1C). Kolmogorov-Smirnoff test of the compulsivity score (*i.e*. averaged number of completed seeking cycles during the last four punishment sessions) confirms that the distribution of the ‘shock-resistant’ population (which completed ≥ 30% of the 10-daily seeking cycles) differs from both ‘shock-sensitive’ and control populations (Fig. S1C: *D* = 1, *P* < 0.001 for both comparisons) while control and ‘shock-sensitive’ groups display comparable distribution (*D* = 0.306, *n.s.*).

**Figure 1.**
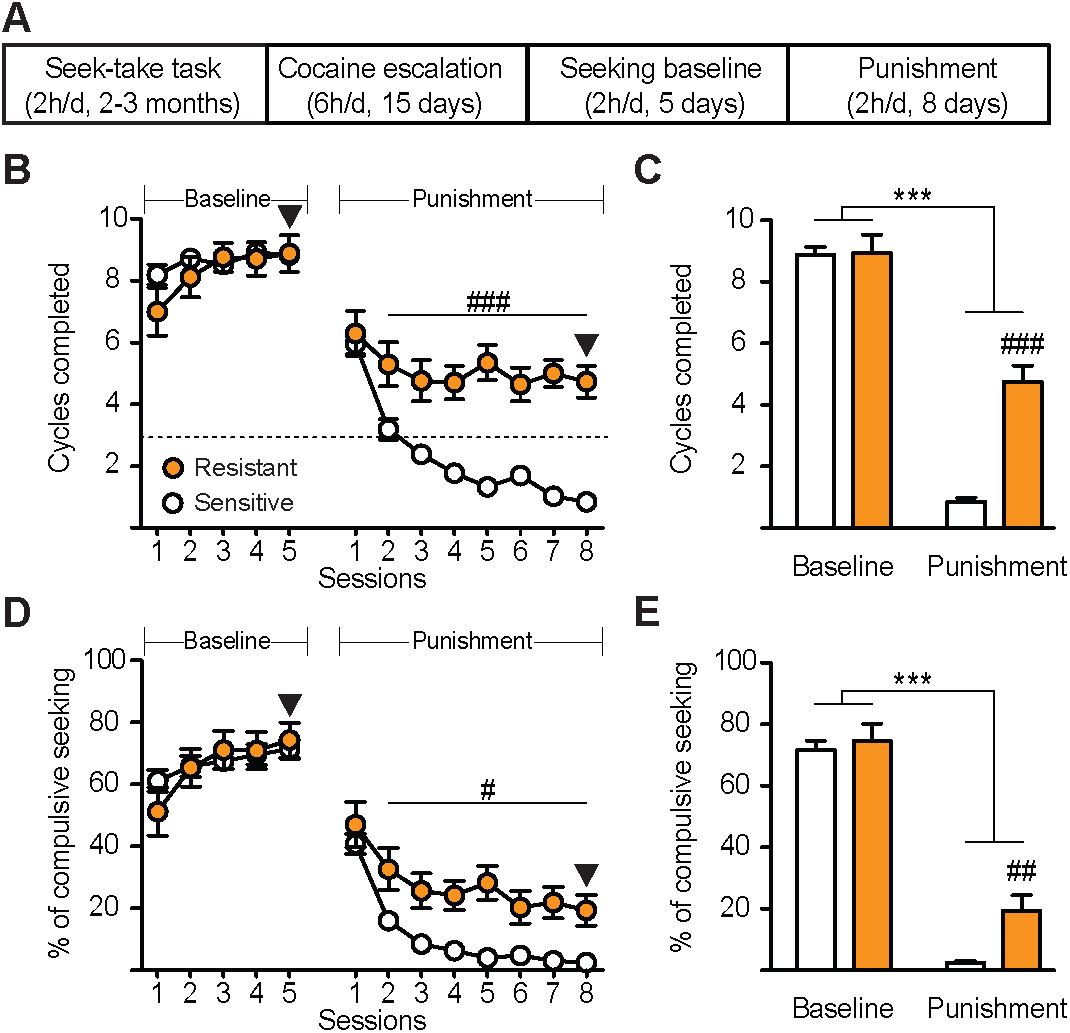
Extended access to cocaine self-administration promotes compulsive cocaine seeking in a subpopulation of rats. (**A**) Experimental time course. Following training on the seek-take task, animals were subjected to cocaine escalation protocol (Fig. S1B). Then, after five days of baseline cocaine seeking, animals were exposed to eight punishment sessions, during which electric foot shock was randomly delivered instead of cocaine in 50% of the trials. (**B**) Punishment contingency reduced the number of seeking cycles completed in all animals (punishment effect: *F*_(*12,612)*_ = 117.2, *P* < 0.0001). While both groups displayed equivalent level of cocaine seeking during baseline (group effect: *F*_(*1,51)*_ = 0.559, *n.s.*), ‘shock-resistant’ rats (*i.e*. ‘compulsive-like’, *n* = 17) completed more seeking cycles than ‘shock-sensitive’ rats (*n* = 36; punishment × group: *F*_(*7,357)*_ = 7.927, *P* < 0.0001; Bonferroni post hoc: ^###^*P* < 0.001 resistant *vs.* sensitive) during punished sessions. Dashed line indicates compulsivity threshold below which animals are considered ‘shock-sensitive’ (see methods). (**C**) Number of seeking cycles completed during the last day of baseline and last day of punishment (arrowheads in B) for both groups (session × group: *F*_(*1,51)*_ = 35.63, *P* < 0.0001; Bonferroni post hoc: \*\*\**P* < 0.001 baseline *vs.* punishment, ^###^*P* < 0.001 resistant *vs.* sensitive). (**D**) Punishment contingency reduced the percentage of compulsive seeking lever presses performed per cycle completed (punishment effect: *F*_(*12,612)*_ = 122.7, *P* < 0.0001). While both groups displayed equivalent level of cocaine seeking during baseline (group effect: *F*_(*1,51)*_ = 0.009, *n.s.*), ‘shock-resistant’ rats performed more compulsive seeking lever presses than ‘shock-sensitive’ rats (group: *F*_(*1,51)*_ = 33.79, *P* < 0.0001; Bonferroni post hoc: ^#^*P* < 0.05 resistant *vs.* sensitive) during punished sessions. (**E**) Percentages of compulsive seeking lever presses performed per cycle completed during the last day of baseline and last day of punishment (arrowheads in **D**) for both groups (session × group: *F*_(*1,51)*_ = 4.184, *P* < 0.05; Bonferroni post hoc: \*\*\**P* < 0.001 baseline *vs.* punishment, ^##^*P* < 0.01 resistant *vs.* sensitive). Line and bar graphs indicate mean ± s.e.m.

As previously reported in tasks for which either drug seeking or drug taking were punished (*4*–*6*, *8*, *10*), punishment contingency diminished the seeking behavior in all animals (Fig. 1B: punishment effect: *F*_(*12,612)*_ = 117.2, *P* < 0.0001). However, ‘shock-resistant’ animals kept seeking cocaine, while the other individuals quickly suppressed this behavior (Fig. 1B: punishment × group: *F*_*(7,357)*_ = 7.927, *P* < 0.0001; Fig. 1C: punishment × group: *F*_(*1,51*)_ = 35.63, *P* < 0.0001). The ‘shock-resistant’ animals performed more compulsive seeking lever presses (*i.e.* presses unnecessary to complete a cycle, see methods) than the ‘shock-sensitive’ rats (Fig. 1D: punishment effect: *F*_*(7,357)*_ = 31.71, *P* < 0.0001, group effect: *F*_*(1,51)*_ = 33.79, *P* < 0.0001; Fig. 1E: punishment × group: *F*_(*1,51*)_ = 4.184, *P* = 0.046).

During seeking sessions, animals were independently given the opportunity to nosepoke for an unpunished sucrose reward. Importantly, sucrose seeking was not altered during punished cocaine seeking sessions in both ‘shock-sensitive’ and ‘shock-resistant’ subpopulations (Fig. S1D: group effect: *F*_*(1,51)*_ = 0.0836, *n.s.*), indicating that the effect of punishment was specific to the punished seeking response and did not reflect a general suppression of responding.

### Frequency-dependent effects of STN DBS on compulsive-like cocaine seeking

Since high frequency stimulation of the STN has beneficial effects on drug-induced addiction-like behaviors (*19*, *20*, *22*), we tested the ability of 130-Hz DBS to reduce compulsive-like cocaine seeking. Following the first eight punishment sessions and characterization of their compulsive status (Fig. 1B), some ‘shock-resistant’ (n = 12) and ‘shock-sensitive’ animals (n = 14) were subjected to 130-Hz STN DBS (*i.e*. ON) during five sessions followed by five punishment sessions with no stimulation (*i*.*e*. OFF) (Fig. S2A). Unexpectedly, 130-Hz STN DBS acutely worsened cocaine compulsive seeking of ‘shock-resistant’ rats: it increased the number of cycles completed on the first two days of stimulation (Fig. S2B: DBS effect: *F*_(*14,336)*_ = 1.764, *P* = 0.0427) as well as the compulsive seeking lever presses on the second day (Fig. S2D: DBS effect: *F*_(*14,336)*_ = 1.718, *P* = 0.0505). However, it did not last throughout the five ON sessions (Fig. S2C: DBS effect: *F*_(*2,48)*_ = 2.489, *n.s.*; Fig. S2E: DBS effect: *F*_(*2,48)*_ = 2.973, *n.s.*). No changes were observed in the seeking behavior of ‘shock-sensitive’ rats during the ON- and OFF-sessions (Fig. S2B: DBS × group effect: *F*_(*14,336)*_ = 0.911, *n.s.*; Fig. S2D: DBS × group effect: *F*_(*14,336)*_ = 0.913, *n.s.)*.

Given these results and since DBS has frequency-dependent effects on drug-induced pathological behaviors in the ventral striatum (*25*, *26*)), we then exposed these animals to STN low frequency DBS (at 30-Hz, as previously tested on motor effects (*27*)) during five punished-seeking sessions followed by five OFF sessions (Fig. 2A). Importantly, to exclude a possible treatment-order effect, half of the animals were first stimulated with 130-Hz DBS and then 30-Hz, and inversely for the other half. 30-Hz STN DBS progressively decreased pathological cocaine seeking throughout the ON-sessions since it reduced both the number of seeking cycles completed by ‘shock-resistant’ animals (Fig. 2B: DBS effect: *F*_(*14,336)*_ = 3.494, *P* < 0.0001) and the number of compulsive seeking lever presses (Fig. 2D: DBS effect: *F*_(*14,336)*_ = 2.6, *P* = 0.0014). These effects also persisted during the OFF-sessions (Fig 2C: *F*_(*2,48)*_ = 9.416, *P* = 0.0004; Fig. 2E: DBS effect: *F*_(*2,48)*_ = 6.168, *P* = 0.0041). Post hoc comparisons between ON sessions in ‘shock-resistant’ animals showed that 30-Hz stimulation reduces both the number of completed cycles (Fig. S2F: *t*_11_ = 3.576, *P* = 0.0043) and compulsive seeking lever presses (Fig. S2G: *t*_11_ = 2.414, *P* = 0.0344) performed during 130-Hz sessions.

**Figure 2.**
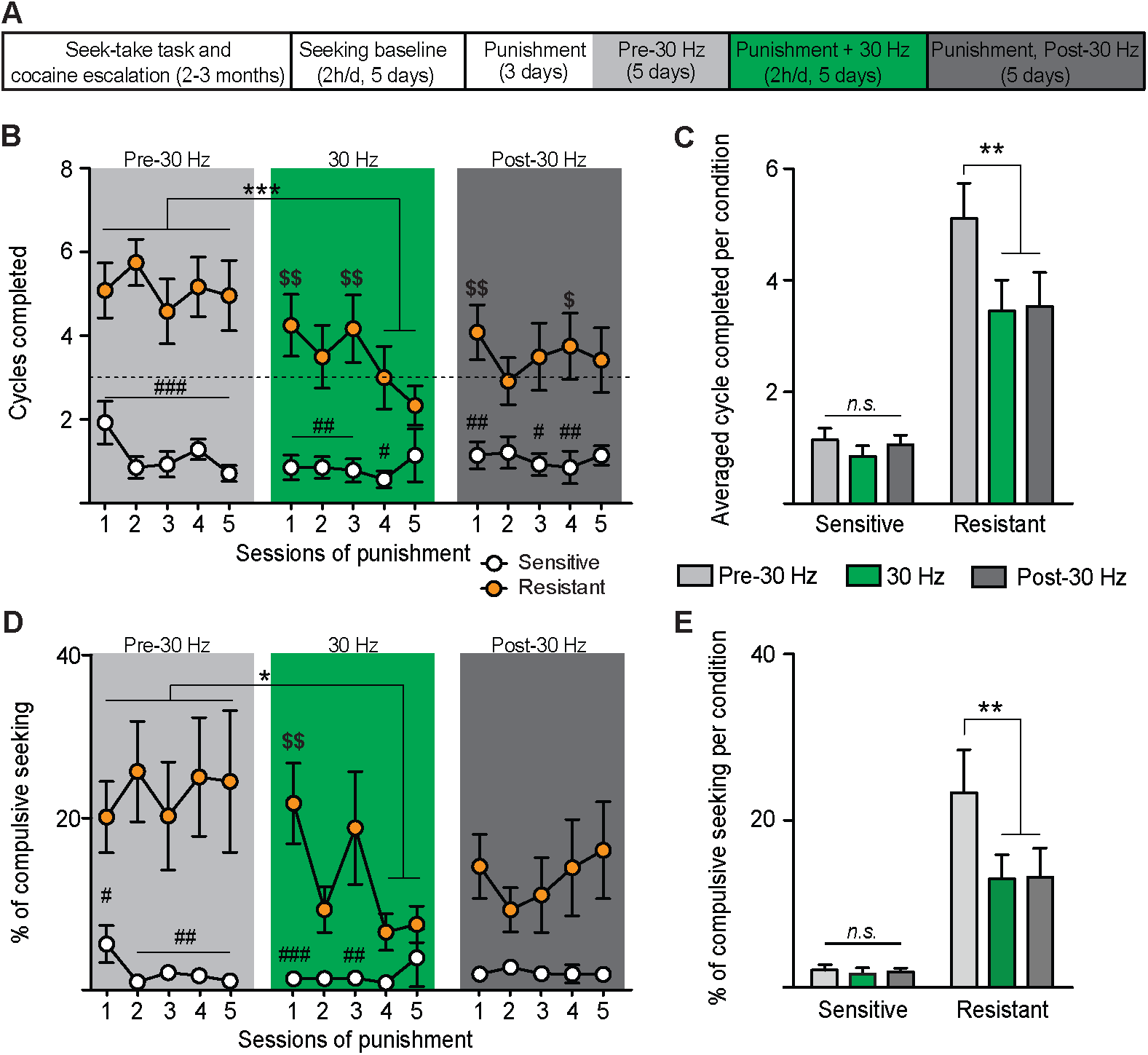
30-Hz STN DBS reduces compulsive-like cocaine seeking during punishment. (**A**) Experimental time course. After characterization of their compulsive status during the baseline-punishment sequence, animals were subjected to five days of 30-Hz STN DBS followed by 5 days with no DBS. (**B**) 30-Hz STN DBS decreased the number of seeking cycles completed by ‘shock-resistant’ rats (orange dots; *n* = 12, DBS effect: *F*_(*14,336)*_ = 3.494, *P* < 0.0001), but had no effect in ‘shock-sensitive’ animals (white dots; *n* = 14, group effect, *F*_(*1,24)*_ = 33.67, *P* < 0.0001; DBS × group effect: *F*_(*14,336)*_ = 2.848, *P* < 0.001; Bonferroni post hoc: \*\*\**P* < 0.001; ^$^*P* < 0.05, ^$$^*P* < 0.01 *vs*. last session of 30-Hz DBS; ^#^*P* < 0.05, ^##^*P* < 0.01, ^###^*P* < 0.001, resistant *vs*. sensitive). Dashed line indicates compulsivity threshold below which animals are considered ‘shock-sensitive’. (**C**) Averaged (5-session block) number of cycles completed by ‘shock-sensitive’ (left) and ‘-resistant’ (right) rats before (pale grey), during (green) and after (dark grey) 30-Hz STN DBS (group effect: *F*_(*1,24)*_ = 33.67, *P* < 0.0001; DBS effect: *F*_(*2,48*)_ = 9.416, *P* < 0.001; DBS × group effect: *F*_(*2,48)*_ = 5.75, *P* < 0.01; Bonferroni post hoc: \*\**P* < 0.01). (**D**) 30-Hz STN DBS decreased the percentage of compulsive seeking lever presses performed by ‘shock-resistant’ rats (DBS effect: *F*_(*14,336)*_ = 2.6, *P* < 0.01), but had no effect in ‘shock-sensitive’ animals (group effect, *F*_(*1,24)*_ = 22.32, *P* < 0.0001; DBS × group effect: *F*_(*14,336)*_ = 2.88, *P* < 0.001; Bonferroni post hoc: \**P* < 0.05; ^$$^*P*<0.01 *vs*. last session of 30-Hz DBS; ^#^*P* < 0.05, ^##^*P* < 0.01, ^###^*P* < 0.001, resistant *vs*. sensitive). (**E**) Averaged (5-session block) percentage of compulsive seeking lever presses performed by ‘shock-sensitive’ (left) and ‘-resistant’ (right) rats before (pale grey), during (green) and after (dark grey) 30-Hz STN DBS (group effect: *F*_(*1,24)*_ = 22.32, *P* < 0.0001; DBS effect: *F*_(*2,48)*_ = 6.168, *P* < 0.01; DBS × group effect: *F*_(*2,48)*_ = 5.389, *P* < 0.01; Bonferroni post hoc: \*\**P* < 0.01).

In the ‘shock-sensitive’ rats, 30-Hz STN DBS affected neither their number of completed cycles (Fig. 2B: group effect, *F*_(*1,24)*_ = 33.67, *P* < 0.0001; DBS × group effect: *F*_(*14,336)*_ = 2.848, *P* = 0.0005; Fig 2C: group effect: *F*_(*1,24)*_ = 33.67, *P* < 0.0001; DBS × group effect: *F*_(*2,48)*_ = 5.75, *P* = 0.0058) nor their compulsive seeking lever presses (Fig. 2D: group effect: *F*_(*1,24)*_ = 22.32, *P* < 0.0001; DBS × group effect: *F*_(*14,336)*_ = 2.88, *P* = 0.0004; Fig. 2E: group effect: *F*_(*1,48)*_ = 22.32, *P*<0.0001; DBS × group effect: *F*_(*2,48)*_ = 5.389, *P* = 0.0077). Post hoc comparisons between ON sessions showed 30-Hz DBS slightly reduced the number of cycles completed by the ‘shock-sensitive’ population during the 130-Hz stimulation (Fig. S2F: *t*_13_ = 2.268, *P* =0.0344) and tended to diminish their compulsive seeking lever presses (Fig. S2G: *t*_13_ = 2.115, *P* = 0.0544).

Of note, 30-Hz STN DBS had no noticeable side effects: it did not promote compensatory sucrose seeking (Fig. S3A: DBS effect for ‘shock-sensitive’ animals; *F*_(*5,65)*_ = 0.573, *n.s.*; Fig. S3B: DBS effect for ‘shock-resistant’ animals; *F*_(*5,55)*_ = 2.099, *n.s.*) and did not affect animals’ basal locomotor activity nor cocaine-induced hyperlocomotion (Fig. S4A: *n* = 6, *F*_(*3,15)*_ = 10.19, *P* = 0.0007). The STN has been recently shown to be involved in pain processing (*28*), suggesting that STN DBS could affect sensitivity to the electric foot shock by modifying peripheral pain sensitivity. We found, however, that five consecutive sessions of high or low frequency STN DBS in naïve rats (n = 9) did not affect their latency to react on a hot plate (Fig. S5: DBS effect: *F*_(*2,12)*_ = 2.034, *n.s.*), ruling out a possible reduced sensitivity to punishment in the stimulated animals.

### STN low frequency oscillations predict compulsive-like cocaine seeking

One major challenge in the addiction research field is to identify vulnerable individuals before they transition to harmful pattern of drug consumption. However, here, ‘shock-sensitive’ and ‘shock-resistant’ rats could not be behaviorally differentiated before the exposure to the punishment. The number of training sessions on the seek-take task was equivalent in both populations (Fig. S6A: *r* = 0.1479, *n.s.*), as well as their seeking performance before being subjected to cocaine escalation, in terms of seeking lever presses (Fig. S6B: group effect: *F*_*(1,51)*_ = 2.512, *n.s.*), cycles completed (Fig. S6C: group effect: *F*_*(1,51)*_ = 2.278, *n.s.*) and compulsive seeking lever presses (Fig. S6D: group effect: *F*_*(1,51)*_ = 2.844, *n.s.*). Likewise after escalation, during which both populations displayed similar cocaine intake (Fig. S1B: group effect: *F*_(*1,51)*_ = 0.4484, *n.s.*) ‘shock-sensitive’ and shock-resistant’ animals exhibited comparable performances during baseline seeking sessions, in terms of seeking lever presses (Fig. S6E: group effect: *F*_(*1,51)*_ = 1.241, *n.s.*), cycles completed (Fig. 1B: group effect: *F*_(*1,51)*_ = 0.559, *n.s.*), compulsive seeking lever presses (Fig. 1D: group effect: *F*_(*1,51)*_ = 0.009, *n.s.*) and sucrose seeking (Fig. S1D: group effect: *F*_(*1,51)*_ = 0.7797, *n.s.*). Also, there was no correlation between animal’s compulsivity score and compulsive seeking lever presses (Fig. S6F: *r* = 0.0599, *n.s.*) or seeking lever presses (Fig. S6F: *r* = 0.0911, *n.s.*) before exposure to punishment contingency.

Our data indicate that compulsive-like cocaine seeking is only observed in some animals that had previously escalated their cocaine intake (see Figs. 1B and S1A,C). We previously showed that cocaine escalation induces pathological oscillations in the STN, compared to activities recorded during short-access sessions (*22*). Thus, one may hypothesize that changes in STN oscillatory activity would predict the rat’s compulsivity status. Therefore, we recorded STN LFPs in some animals (from Fig. 1), through DBS electrodes, during cocaine escalation, when their compulsivity status was unknown. STN activity was monitored during 15-min before, and after, the 6-h session of cocaine intake on days 1, 4, 8, 12 and 15 of the escalation procedure (Fig. 3A: session effect: *F*_*(14,210)*_ = 6.675, *P* < 0.0001; group effect: *F*_*(1,15)*_ = 0.813, *n.s.*). Following the punishment sessions and the characterization of each rats’ compulsivity status (Fig. 3B: punishment × group: *F*_*(7,105)*_ = 3.947, *P* = 0.0007), post hoc analysis of STN LFPs power spectrum revealed two distinct patterns of activity during the development of cocaine escalation (Fig. S7A,B). Compared to basal oscillatory activity recorded on the first day of escalation, ‘shock-resistant’ animals recorded (n = 5) exhibited a progressive increase in STN low (6-40 Hz), but not high (65-90Hz), frequency oscillation powers during LFPs baseline recordings (*i.e.* before cocaine access) that was not observed in the ‘shock-sensitive’ rats recorded (n = 12) (Fig. 3C). Band-specific analysis revealed significant power increases of STN oscillations in alpha/theta (6-13Hz, Fig. 3D: session effect: *F*_*(4,60)*_ = 5.481, *P* = 0.0008) and low/high beta bands (14-40Hz, Fig. 3E: session effect: *F*_*(4,60)*_ = 4.119, *P* = 0.0051), but not in the gamma band (65-90Hz, Fig. 3F: session effect: *F*_*(4,60)*_ = 0.9093, *n.s*.) during escalation of cocaine intake, only in ‘shock-resistant’ rats (group effect: *F*_*(1,15)*_ = 5.454 and 6.035, *P* = 0.034 and 0.027 for alpha/theta and beta bands, respectively). Of note, both populations exhibited comparable levels of activity on the first day of escalation (Fig. 3D-F). In line with the ability of dopaminergic agonist treatment to reduce STN pathological oscillations in both PD monkeys and patients (*14*, *15*, *29*), these pathological increases were no longer present immediately after cocaine self-administration (Fig. 3D-E: session × group effect: *F*_(*9,135)*_ = 4.059 and 3.514, *P* = 0.0001 and 0.0006 for alpha/theta and beta bands, respectively; Fig. S7B), thereby suggesting a dopamine-dependent mechanism sustaining the onset of these pathological oscillations. We also found that the baseline oscillatory activity before the last escalation session and the animal’s compulsivity score was positively correlated for alpha/theta (Fig. S7C: *r* = 0.6048, *P* = 0.01) and beta bands (*r* = 0.5127, *P* = 0.04) but not for the gamma band (*r* = 0.1869, *n.s.*). Finally, analysis of basal level of oscillation power changes of ‘shock-resistant’ rats between the first and the last cocaine escalation sessions indicates that oscillations around 8-Hz were the most dramatically affected at the end of escalation (173.78 ± 49.82% increase, Fig. S7D: session effect: *F*_(*1,252)*_ = 286.9, *P* < 0.001; session × frequency effect: *F*_(*62,252)*_ = 1.548, *P* = 0.01). These animals also displayed an increase around 18-Hz (141.94 ± 59.95% increase).

**Figure 3.**
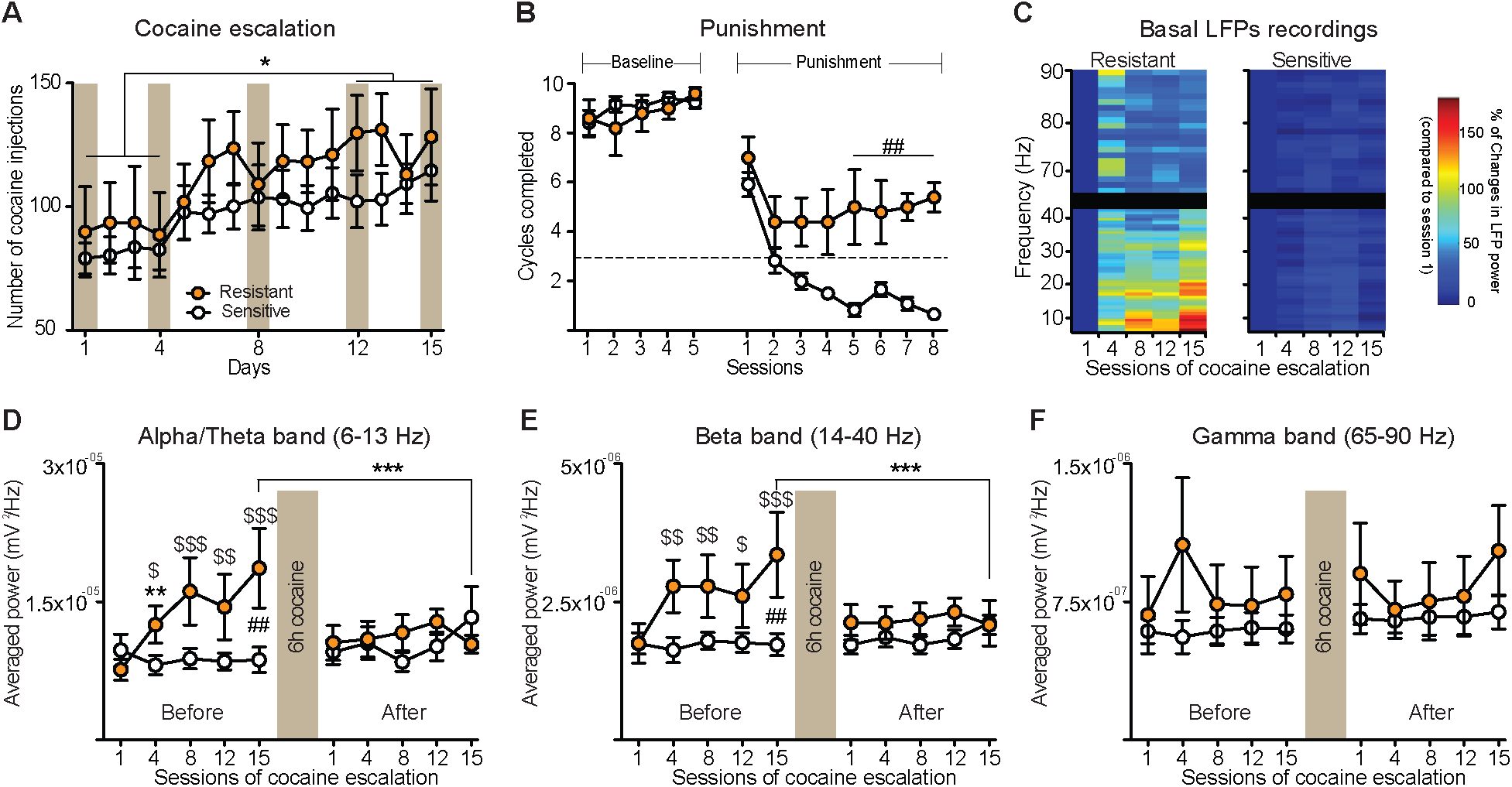
Compulsive rats exhibit pathological STN low frequency oscillations during cocaine escalation. (**A**) Future‘shock-sensitive’ (*n* = 12) and ‘shock-resistant’ (*n* = 5) rats exhibited similar drug intake during cocaine escalation (session effect: *F*_(*14,210*)_ = 6.675, *P* < 0.0001; group effect: *F*_*(1,15)*_ = 0.813, *n.s.*; Bonferroni post hoc: \**P* < 0.05). Brown rectangles indicate LFPs recording sessions. (**B**) Punishment contingency reduced the number of seeking cycles completed in all animals (session effect: *F*_(*12,180)*_ = 66.21, *P* < 0.0001). While both groups displayed equivalent level of cocaine seeking during baseline (group effect: *F*_(*1,15)*_ = 0.2039, *n.s.*), ‘shock-resistant’ rats completed more seeking cycles than ‘shock-sensitive’ rats (punishment × group: *F*_*(7,105)*_ = 3.947, *P* < 0.001; Bonferroni post hoc: ^##^*P* < 0.01 resistant *vs.* sensitive) during punished sessions. Dashed line indicates compulsivity threshold below which animals are considered ‘shock-sensitive’. (**C**), Session-frequency power spectrum showing basal (*i.e.* before cocaine) LFPs power changes normalized (in %) to session 1 in ‘shock-resistant’ (left) and ‘shock-sensitive’ (right) animals. (**D-F**) Quantifications of LFPs power before and after 6h of cocaine access. During baseline recordings, ‘shock-resistant’ rats showed a progressive power increase in alpha/theta (**D**, session effect: *F*_(*4,60)*_ = 5.481, *P* = 0.0008) and beta (**E**, session effect: *F*_(*4,60)*_ = 4.119, *P* = 0.0051; Bonferroni post hoc: ^$^*P* < 0.05, ^$$^*P* < 0.01, ^$$$^*P* < 0.001 vs. session 1; \*\**P* < 0.01 vs. session 15), but not in gamma band (**F**, session effect: *F*_*(4,60)*_ = 0.909, *n.s.*). No increase was observed in ‘shock-sensitive’ animals (group effect: **D**, *F*_(*1,15)*_ = 5.454, *P* = 0.034; **E**, *F*_(*1,15)*_ = 6.035, *P* = 0.027; Bonferroni post hoc: ^##^*P* < 0.01 resistant *vs.* sensitive). Cocaine consumption reduces increased power on session 15 in both alpha/theta and beta bands (session × group effect: **D**, *F*_(*9,135)*_ = 4.059, *P* = 0.0001; **E**, *F*_(*9,135)*_ = 3.514, *P* < 0.001; Bonferroni post hoc: \*\*\**P* < 0.001) but has no effect in the gamma band (**F**, session × group effect: *F*_(*9,135)*_ = 1.393, *n.s.*). Line graphs indicate mean ± s.e.m.

### Causal contribution of STN low frequency oscillations to the onset of compulsive-like cocaine seeking

To test the causal role of increased STN low frequency oscillations on the emergence of compulsive-like cocaine seeking, we first wondered whether an 8-Hz DBS could induce STN low-frequency oscillatory activity. Naïve rats (n = 4) were implanted with DBS electrodes in the STN. After recovery, they were subjected to 8-Hz STN DBS (6h/day) during 7 days. LFPs were recorded before and after the first 6-h stimulation session and after the last DBS session on day 7. Given that (1) these animals were not subjected to any behavioral procedures and (2) each unilateral DBS electrode has its own electrical reference (see methods), left or right STN (n = 5; 3 were discarded for misplacement of the electrode) were analyzed independently. 6h of 8-Hz STN DBS markedly increased STN low, but not high, frequency oscillatory activity (Fig. S8A: session effect: *F*_(*2,1112)*_ = 128.1, *P* < 0.0001; session × frequency *F*_(*276,1112)*_ = 3.461, *P* < 0.0001), notably in the theta band (6-13 Hz), and at a lesser degree in the low beta band (14-18 Hz). No further increase was observed after 7 days of stimulation. Band-specific analysis confirmed that an 8-Hz DBS can induce low frequency oscillatory activity, especially in the alpha/theta band (Fig. 8B: *F*_(*2,8)*_ = 4.465, *P* = 0.0499; beta band: *F*_(*2,8)*_ = 2.846, *n.s.*; gamma band: *F*_(*2,8)*_ = 1.201, *n.s.*).

We next tested whether applying alpha/theta (8 Hz) stimulation in the STN of ‘shock-sensitive’ animals during a second cocaine escalation (escalation-2), thus mimicking abnormal very low frequency oscillations, could change their compulsivity status upon re-exposure to the punishment protocol (punishment-2) (Fig. 4A). Following the first escalation-punishment sequence (escalation-1 and punishment-1) and characterization of their compulsivity profile, some ‘shock-sensitive’ rats (*N* = 13; Fig. 4B: punishment effect: *F*_(*7,70)*_ = 29.61, *P* < 0.0001; group effect: *F*_(*2,10)*_ = 0.2029, *n.s.*) were subjected to escalation-2, during which some were stimulated within the STN at 8-Hz (*n* = 5) or 70-Hz (a frequency at which no significant changes were observed in STN oscillation power on the last session of cocaine escalation, *n* = 3, see Fig. S7D), while the control rats (*n* = 5) were not stimulated. STN DBS at 8-Hz or 70-Hz had no effect on cocaine intake during escalation-2 (Fig. 4C: session effect: *F*_(*13,130)*_ = 5.302, *P* < 0.0001; group effect: *F*_(*2,10)*_ = 0.1681, *n.s.*), indicating that only STN 130-Hz DBS can reduce re-escalation of drug intake (*20*, *22*)), and did not change the rats’ pattern of cocaine consumption observed during escalation-1 (Fig. S9A: session effect: *F*_(*14,140)*_ = 3.136, *P* = 0.0003.; group effect: *F*_(*2,10)*_ = 0.3937, *n.s.*; Fig. S9B: session effect: *F*_(*13,140)*_ = 0.601, *n.s.*; group effect: *F*_(*2,140)*_ = 1.079, *n.s.*).

**Figure 4.**
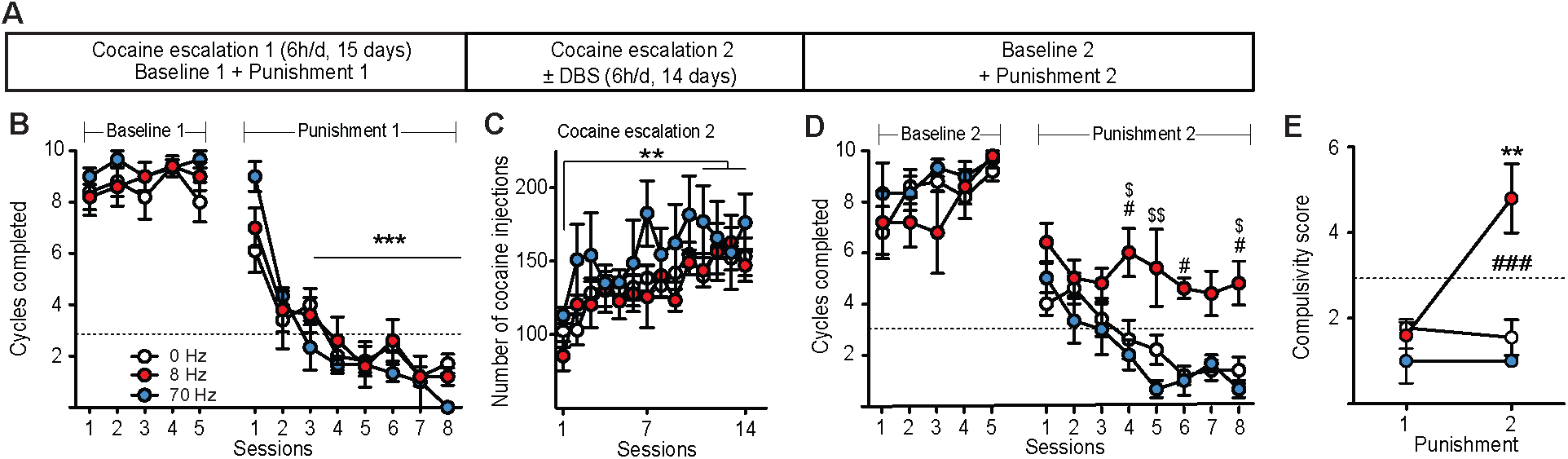
STN 8-Hz DBS triggers cocaine compulsive seeking in ‘shock-sensitive’ rats. (**A**) Experimental time course. Following cocaine escalation and punishment protocols and characterization of the compulsive status of the animals, shock-sensitive rats were subjected to a second escalation procedure during which some of the animals received STN DBS at 8-Hz (red dots, *n* = 5) or at 70 Hz (blue dots, *n* = 3), while the others remained OFF stimulation (white dots, *n* = 5). They were then all tested a second time in the punishment paradigm. (**B**) Punishment delivery suppressed cocaine seeking in all groups (punishment effect: *F*_(*12,120)*_ = 89.94, *P* < 0.0001; group effect: *F*_(*2,10)*_ = 0.168, *n.s.*; Bonferroni post hoc: \*\*\**P* < 0.001, *vs.* baseline). All groups displayed equivalent level of cocaine seeking during baseline 1 (group effect: *F*_(*2,10)*_ = 0.449, *n.s.*) and punishment 1 (group effect: *F*_(*2,70)*_ = 0.2029, *n.s.*). (**C**) STN DBS during cocaine escalation 2 did not alter escalation of cocaine intake (session effect: *F*_(*13,130)*_ = 5.302, *P* < 0.0001; group effect: *F*_(*2,10)*_ = 2.142, *n.s.*; Bonferroni post hoc: \*\**P* < 0.01). (**D**) Punishment contingency during punishment 2 reduced the number of seeking cycles completed in all groups (punishment effect: *F*_(*12,120)*_ = 40.94, *P* < 0.0001). While all groups displayed equivalent level of cocaine seeking during baseline 2 (group effect: *F*_(*2,10)*_ = 0.529, *n.s.*), 8-Hz-stimulated rats (red dots) completed more seeking cycles than 0 Hz- (black dots) and 70Hz-stimulated rats (blue dots) (punishment effect: *F*_(*7,70)*_ = 9.108, *P* < 0.0001; group effect: *F*_(2*,10)*_ = 10.48, *P* = 0.0035; Bonferroni post hoc: ^#^*P* < 0.05, 8-Hz *vs.* 0 Hz; ^$^*P* < 0.05, ^$$^*P* < 0.01, 8-Hz *vs.* 70 Hz) during punishment 2. (**E**) Compulsivity score before and after cocaine escalation 2 (before/after × group effect: *F*_(*2,10)*_ = 7.705, *P* = 0.0094; Bonferroni post hoc: \*\**P* < 0.001, before *vs.* after, ^###^*P* < 0.001, 8-Hz *vs*. 0 / 70 Hz). Dashed lines indicate compulsivity threshold below which animals are considered shock-sensitive. Line graphs indicate mean ± s.e.m.

Following escalation-2, the number of cycles completed during baseline-2 was similar between groups (Fig. 4D: group effect: *F*_(*2,10)*_ = 0.529, *n.s*) and did not differ from baseline-1 (Fig. S9C: group effect: *F*_(*2,10)*_ = 0.3723, *n.s*.). Likewise, all groups exhibited comparable level of compulsive seeking lever presses during the last session of baseline-2, which was not different from baseline-1 (Fig. S9D: session effect: *F*_(*1,10)*_ = 1.791, *n.s.*, group effect: *F*_(*2,10)*_ = 3.276, *n.s.*). In contrast, when re-exposed to the foot shock contingency, the 8-Hz-stimulated rats displayed ‘shock-resistant’-like behaviors. Indeed, they completed more seeking cycles than during punishment-1 (Fig. S9C: session effect: *F*_(*7,70)*_ = 9.131, *P* < 0.0001; group effect: *F*_(*2,10)*_ 9.532, *P* = 0.0048) and than other groups during punishment-2 (Fig. 4D: session effect: *F*_(*7,70)*_ = 9.108, *P* < 0.0001; group effect: *F*_(2*,10*)_ = 10.48, *P* = 0.0035), which quickly stopped seeking cocaine. Accordingly, their compulsivity score was increased (Fig. 4E: before/after × group effect: *F*_(*2,10)*_ = 7.705, *P* = 0.0094) and they performed more compulsive seeking lever presses on the last session of punishment-2, compared to punishment-1 (Fig. S9D: session effect: *F*_(*1,10)*_ = 6.342, *P* = 0.03). Importantly, sucrose seeking level was not altered in any group, compared to baseline-1 and punishment-1 (Fig. S9E: session effect: *F*_(*12,130)*_ = 0.8742, *n.s.*; group effect: *F*_(*2,130)*_ = 0.9482, *n.s.*), indicating that 8-Hz STN DBS did not promote a general increase in reward seeking behaviors. To rule out a possible effect of STN DBS on pain perception that would had promoted compulsive-like cocaine seeking, we showed that a similar 8-Hz treatment (6 h/day for 14 days) applied to other rats (*n* = 4) did not affect peripheral pain perception on a hot plate (Fig. S10A: stimulation effect *F*_(*2,6)*_ = 3.34, *n.s.*).

## Discussion

Decades of dedicated research have profoundly expanded our knowledge regarding the short- and long-term consequences of drug consumption, at both the cellular and behavioral levels, in rodents and humans. However, specific biomarkers predictive of the pathology are still lacking to help detecting vulnerable individuals and designing effective personalized treatments that would improve long-term recovery of drug addicts (*30*).

Using a rodent model of compulsive-like cocaine seeking behavior, which only emerges in some animals that had previously escalated their cocaine intake, we show that abnormal oscillatory activity within the STN predicts the compulsivity status of animals. Specifically, a pathological increase in STN alpha/theta oscillatory activity during cocaine escalation occurs only in animals that were subsequently seeking cocaine despite the foot-shock contingency during the punishment protocol. These abnormal increases in low frequency power observed only in the future ‘resistant’ animals could be a marker of two major phenomena: first, the abnormal increases might represent an indicator of the craving resulting from the daily 18h-abstinence period experienced by animals during the escalation protocol or second, these abnormal increases could predict the rat’s seeking phenotype. Since the abstinence was the same for all rats and no changes in oscillatory activity were detected in future ‘sensitive’ animals, the pathological increases in STN low frequency oscillation power may represent a specific predictive biomarker of compulsivity or vulnerability to compulsively seek drug.

The lack of increase in STN oscillatory activity of ‘shock-sensitive’ animals may appear in contradiction with our former study, which showed that changes in baseline STN neuronal activity correlates with the loss of control during cocaine escalation (*22*). Several experimental considerations have to be elaborated. One major difference between our two studies concerns the animals’ history of cocaine intake. We showed previously that in rats naïve to cocaine, a short access to cocaine (2 h per day) for ten days induces a slight increase in STN low frequency oscillations during baseline recordings and this increase is further enhanced during escalation of cocaine intake (*22*). Thus, the change in basal low frequency activities during escalation was assumed to be related to STN activity during cocaine short access sessions. In contrast, here, rats have been trained in the seeking/taking task during several months, allowing them to self-administer cocaine daily, but STN activity was only recorded from the beginning of the cocaine escalation protocol. Therefore, the reference for the measurement of the power change is totally different (*i.e.* the baseline is different). Overall, these data confirm the critical role of STN activity, especially in its low frequencies, in the transition to- and the expression of addictive-like behaviors.

While cocaine re-escalation did not promote compulsive-like seeking by itself in control ‘shock-sensitive’ animals, 8-Hz-stimulated rats exhibit the compulsive-like trait upon re-exposure to the punishment, thereby confirming the causal predictive nature of STN low frequency activity to compulsive-like seeking. Several studies have outlined the contribution of STN low frequency oscillations (alpha/theta band) in motor, emotional and cognitive processes in PD and OCD patients. For instance, they correlate with levodopa-induced dyskinesia and impulse control disorders in PD patients (*18*, *31*)). Also, decision-making in a high-conflict task has been shown to increase STN theta oscillations (*32*, *33*)), which are coordinated with theta activity of the prefrontal cortex during conflict detection (*34*, *35*)). Also, the severity of compulsive, but not obsessive, symptoms in OCD patients is correlated to STN oscillatory activity within the theta band (*16*), which is bi-directionally modulated by failure or success to inhibit OCD symptoms (*17*). As such, emergence of STN alpha/theta activity in ‘shock-resistant’ rats (or maintaining an 8-Hz activity in ‘shock-sensitive’ animals) during cocaine escalation might thus alter decision-making processes and promote compulsive-like and impulsive-like behaviors that would be fully expressed during the exposure to the punishment, when animals have to face the choice to seek drug despite negative consequences.

To date, pharmacological and/or behavioral therapies have failed to produce consistent and long-lasting beneficial effects for humans suffering from cocaine addiction. The data presented here demonstrate the ability of STN 30-Hz, but not 130-Hz, DBS to reduce compulsive-like cocaine seeking behavior in ‘shock-resistant’ rats. Of importance, such treatment did not affect the seeking behavior of ‘shock-sensitive’ rats and did not engender noticeable side effects. Given its efficiency and safety for human neurodegenerative and psychiatric disorders (*36*, *37*)), our present and previous work (*19*, *20*, *22*) strongly emphasize the therapeutic potential of STN DBS for the treatment of addiction. However, in order to avoid deleterious effects, as previously noted in PD patients (*38*), the frequency seems to be critical. Indeed, we show here that the effect observed are frequency-dependent: 8-Hz STN DBS, which does not affect cocaine re-escalation can trigger compulsive cocaine seeking in non-compulsive animals, while in contrast 130-Hz STN DBS reduces cocaine escalation and relapse (*22*), but can transiently increase pathological seeking of compulsive rats. Thus, the development of STN DBS-based personalized interventions, in which specific frequencies could be applied at precise stages of drug intoxication, may be of critical importance to efficiently normalize pathological seeking and consummatory behaviors toward a more recreational/controlled pattern of use. Such strategy has recently been outlined for DBS use in PD (*39*).

Here, we uncovered a STN DBS bi-directional control over compulsive cocaine seeking (Fig. S9), which is frequency (8-Hz *vs.* 30-Hz)- and addiction stage (escalation *vs*. punished seeking)-dependent. Mechanisms sustaining these beneficial and deleterious effects remain to be elucidated. In rodents and non-human primates, low frequency stimulations (~20 Hz) can drive STN neurons activity both *in vitro* and *in vivo* (*40*, *41*)), which would then alters STN output structure activities to promote or worsen PD-like motor symptoms (*38*, *42*)). Alternatively, antidromic activation of the hyperdirect pathway, which monosynaptically connects cortical areas to the STN in mammals (*43*–*45*), has been proposed to mediate beneficial effects of STN DBS in PD (*46*, *47*)). Activity of the prefrontal cortex, which is involved in decision-making and reward seeking (*48*), is profoundly affected in human drug users (*49*). Moreover, following cocaine escalation and exposure to a punished cocaine seeking paradigm, prefrontal neurons of ‘shock-resistant’ animals are more hypoactive than those of ‘shock-sensitive’ (*4*). Also, STN low, compared to high, frequency stimulations are more efficient to antidromically and accurately drive prefrontal neuron activity (*47*). In such a scenario, 8-Hz STN DBS may thus impose a pathological hypoactivity to prefrontal cortex neurons that would subsequently promotes compulsive-like drug seeking behaviors. Conversely, 30-Hz STN DBS might antidromically ‘re-boost’ prefrontal neuron activity to reduce compulsive-like cocaine seeking, as observed with local low frequency photostimulation (*4*). However, a possible influence of orthodromic activation of STN output structures, thus affecting other pathways involved in compulsive-like behaviors (*e.g.* the orbitofrontal cortex to the dorsal striatum pathway) (*50*) cannot be ruled out.

We highlight the predictive nature of abnormal low-frequency oscillations in the STN during the transition from extended cocaine access to the onset of compulsive-like seeking behaviors. Such a marker may be of critical importance for the development of new diagnostic tools and prevention strategies in humans. Indeed, future investigations aiming at detecting cortical correlates of STN abnormal activity, as explored in PD (*51*), may help to detect vulnerable subjects that could develop pathological drug seeking, in a non-invasive manner (*e.g.* electroencephalography). We also show that the efficient frequency applied in STN depends on the stage of addiction. This could lead to close-loop and adaptive techniques of STN DBS for addiction.

## Materials and Methods

### Study design

The main objective of this study was to investigate the role of STN activity in the onset of compulsive-like seeking behaviors in rats and to test whether altering this activity with DBS could modify this pathological behavior. Animals were thus implanted with DBS electrodes within the STN, thereby allowing recording of activity by measuring LFPs or delivering electrical stimulation at different frequencies. No statistical methods were used to predetermine sample size, but they are comparable to those reported in previous studies (*5*, *19*, *22*). Power analysis was further performed with the G*Power software to confirm that our sample sizes were sufficient to detect reliable changes for critical experiments (power ≥ 80% at a level of confidence p<0.05; values are reported in the statistical table S1). Group assignment was pseudo-randomized for most experiments, as rats were assigned to experimental group in a counterbalanced fashion based on their initial basal cocaine seeking levels. Treatment assignment was randomized between experimental groups. All experiments (except locomotor and pain tests) have been replicated at least twice. Animals were excluded from the study after histological assessments in case of electrodes implantation outside the STN by an experimenter blinded to the experimental conditions.

### Animals

Adult Lister Hooded males (~380 g, Charles River, N = 84) were paired housed, in Plexiglas cages and maintained on an inverted 12h light/dark cycle (light onset at 7 pm) with food and water available *ad libitum*, in a temperature- and humidity-controlled environment. All animal care and use conformed to the French regulation (Decree 2010-118) and were approved by local ethic committee, the University of Aix-Marseille and the Ministry (Saisine #3129.01).

### Electrode design for STN DBS or LFP recordings

The electrodes were made out of Platinum-Iridium wires coated with Teflon (75 μm). Coating was removed over 0.5 mm at the tips and two wires were inserted into a 16 mm stainless steel tubing to form an electrode. Two electrodes, separated by a distance of 4.8 mm (*i.e.* twice the STN laterality), were soldered to an electric connector, allowing connection with both recording and stimulation devices. Electrodes (impedance = 20 kΩ ± 2.25) and connector were subsequently deeply bound, using a custom mold and dental cement. Finally, electrodes were tested with an isolated battery to avoid electrical short circuits.

### Catheter and stereotaxic surgeries

Rats were implanted with a chronically indwelling intravenous catheter, as previously described (*19*, *22*)). Briefly, rats were anesthetized with ketamine (Imalgen, Merial, 100 mg/kg, *s*.*c*.) and medetominine (Domitor, Janssen, 30 mg/kg, *s.c.*) following a preventive long-acting antibiotic treatment (amoxicillin, Duphamox LA, Pfizer, 100 mg/kg, *s*.*c*.). A homemade silicone catheter (0.012-inch inside diameter, 0.025-inch outside diameter, Plastics-One) was inserted and secured into the right jugular vein. The other extremity of the catheter was placed subcutaneously in the mid-scapular region and connected to a guide cannula secured with dental cement. Animals were then placed in a stereotaxic frame (David Kopf apparatus) and maintained under ketamine/medetominine anesthesia. Electrodes were inserted bilaterally in the STN (in mm: −3.7 AP, ±2.4 L from bregma, −8.35 DV from skull surface (*52*), with the incisor bar at −3.3 mm). Four anchoring screws (the one on the right frontal lobe designated as the electric reference allowing LFP recordings in some animals) were fixed into the skull. Electrodes, screws and skull were deeply bounded with dental cement. After surgery, rats were awakened with an injection of atipamezol (Antisedan, Janssen, 0.15 mg/kg *i*.*m*.) and allowed to recover for at least 7 days with *ad libitum* access to food and water. The catheters were daily flushed during the recovery period and just before and after each self-administration session with a saline solution containing heparin (Sanofi, 3 g/l) and enroflorilexine (Baytril, Bayer, 8g/L) to maintain their patency and to prevent infection. Catheters were also regularly tested with propofol (Propovet, Abbott, 10 mg/ml) to confirm their patency.

### Behavioral apparatus

#### Self-administration apparatus

behavioral experiments were performed during the dark phase and took place in standard rat operant chambers (MedAssociates), located in sound-attenuating cubicles, equipped with a house light, two retractable levers, which flanked a sucrose magazine, set 7 cm above the metallic grid floor through which an electric foot shock could be delivered via a generator (MedAssociates). A cue light was positioned 8 cm above each lever. For each rat, one lever was randomly paired with cocaine infusion (taking-lever) while the other one was designated as the seeking-lever. For intravenous drug administration, the stainless-steel guide cannula of the catheter was connected through steel-protected tygon tubing to a swivel (Plastics One) and then an infusion pump (MedAssociates). Data were acquired on a PC running MED-PC IV (MedAssociates). Sessions lasted for 2 h or 6 h (see below for detailed procedures).

#### Locomotor activity apparatus

Locomotor activity was measured as the distance traveled (in meters) in a circular home-made Perspex open field (60 cm diameter). A video tracking system was placed above the open field. Data were acquired by the software Bonsai (Open Ephys), recorded on a PC computer and analyzed offline with Matlab.

#### Hot plate apparatus

Animals were placed on a hot plate analgesia meter (Harvard apparatus) maintained at 52.0 ± 0.5 °C. Rats were continiously observed during the test to detect the first sign of pain (paw licking, rapid movements, escape…) to quickly remove them from the apparatus. Animal’s behavior was recorded by a video tracking system. Data were acquired by the software Bonsai (Open Ephys). Latency to react was quantified offline by an experimenter blind to the DBS treatment.

### Acquisition of cocaine self-administration under the seek-take task schedule

At least one week after surgery, rats began cocaine self-administration training using the seek-take chain schedule, adapted from the previously described procedure (*5*). Self-administration training was divided into four distinct phases: acquisition of the taking response; training on the seek-take chain; extended self-administration and punishment.

### Acquisition of the taking response

In this initial phase, each trial started with the illumination of the house light and the insertion of the taking-lever. One press on the lever, under a fixed ratio schedule (FR-1), resulted in the delivery of a single infusion of cocaine (250 μg/90 μL over 5 s, Coopérative pharmaceutique française). Cocaine infusions were paired with illumination of the cue light (5 s) above the taking lever, retraction of the taking-lever and extinction of the house light. Following a 20 s time out-period, another trial was initiated with the insertion of the taking-lever. Training of the taking response continued, typically for six to eight sessions, until animals reached a stable level of cocaine intake (< 20% changes across 3 consecutive sessions), after which they advanced to the seeking-taking chain schedule.

### Training on the seeking-taking chain schedule

Each cycle started with the illumination of the house light, insertion of the seeking-lever and retraction of the taking-lever. A single press on the seeking-lever resulted in the retraction of the seeking-lever and the insertion of the taking-lever, which activation then triggered cocaine delivery, illumination of the associated cue-light and retraction of the taking-lever and extinction of the house light. Following a 20 s time-out refractory period, another cycle was initiated with the insertion of the seeking-lever.

Once animals reached a stable level (< 20% variation in number of cycles completed across 3 consecutive sessions), a random interval (RI) schedule was introduced into the seeking-link of the chain schedule. Here, the first seeking-lever press initiated a RI schedule of 2 s, which progressively increased to 15, 30, 60 and 120 s during training. Seeking-lever presses within the RI had no programmed consequences, but were still recorded (see below). The first seeking-lever press following the end of the RI lead to the retraction of the seeking-lever and the insertion of the taking-lever, which press triggered the cocaine delivery, paired with the illumination of the associated cue-light as during the training of the taking response, thereby ending the seeking-taking cycle. Each cycle was followed by a 2 min time-out (TO) period, where both levers were retracted, which progressively increases to 4 and 10 min over consecutive training days, before initiation of the next seek-take cycle. Training on each RI-TO schedule persisted until animals displayed less than 20% variation in the numbers of cycle completed. Once an animal achieved stable cocaine seeking behavior at a RI-TO schedule, usually 5-7 days, it was advanced to the next RI-TO schedule.

During these sessions, rats were also trained to nose poke into the sucrose magazine to obtain 0.04 ml of a 20% sucrose solution, which was delivered under a RI schedule, which parameter was progressively increased to 60 s. Sucrose seeking-taking behavior occurred concurrently and independently to the cocaine seeking-taking schedule, thereby allowing us to specifically investigating ‘natural’ reward seeking.

At the end of training under the seeking-taking schedule, animals were allowed to complete up to 10 cocaine cycles and 120 sucrose deliveries in each 2 h session of the RI120-TO10 schedule.

### Extended cocaine self-administration

After reaching the RI120-TO10 schedule criteria of the seeking-taking schedule (< 20% variation in number of cycles completed across 3 consecutive sessions), all animals (except those Fig. S1A, which underwent punishment protocol right after completed RI120-TO10 schedule), were submitted to escalation protocol, in which they were allowed to self-administer cocaine over 6 h for 15 consecutive days to loss control over their cocaine intake (*24*), a cardinal criteria of human drug addiction (*1*). Here, sessions started with the illumination of the house light and the insertion of the taking-lever. As in the training phase, each press on this lever triggered cocaine delivery and the illumination of the associated cue-light, followed by a 20 s TO. Sucrose seeking was unavailable during extended cocaine self-administration sessions. After 15 days, animals were re-subjected to the seek-take RI120-TO10 schedule, as described above, during five days to establish post-escalation cocaine seek-take baseline.

### Punishment

After RI120-TO10 baseline, animals were subjected to 8 daily sessions under the resistance to punishment paradigm (*5*). Here, half of the completed RI120-TO10 cycles resulted in mild foot shock delivery (0.5 mA, 0.5 s) to the animals’ paws: punishment was administered following the first press on the seeking-lever after the RI120 schedule has elapsed, with no access to the taking lever. The other half of the RI120 completed triggered the insertion of the taking lever, which press initiated cocaine delivery and illumination of the associated cue-light.

Seeking cycle’s outcomes (foot-shock or insertion of the cocaine taking lever) were delivered in a pseudorandomized manner, so that difference between both outcomes cannot exceed 2. As such, if the two first seeking cycles lead to two foot shocks, the third cycle will automatically give access to the cocaine taking lever. Thus, depending on their performances on the punished seek take task, animals can receive up to five foot-shocks and five cocaine infusions.

Compulsivity score was defined as the average number of completed cycles (foot-shock received + cocaine deliveries) during the last four sessions of the punishment paradigm. Animals with a compulsivity score ≥ 3 (*i.e.*, completed more than 30% of the 10-daily seeking cycles during the last four sessions of the punishment paradigm) were classified as ‘compulsive’ or ‘shock-resistant’.

To further evaluate compulsive lever press behaviors in rats, we established the percentage of compulsive seeking lever presses per cycle completed, which computes the numbers of total and futile (those unnecessary to complete a cycle) lever presses and the daily seeking performance (*i.e.* number of cycles completed over the 10-daily cycles available) of animals, as follows: %Compulsive lever presses = 100* ([futile lever presses/total lever presses] * [cycles completed/total cycles]).

### Spontaneous- and cocaine-induced locomotor activity measurement

At least 2 months after completing the experiment, *n* = 6 animals were tested on locomotor activity effect of STN DBS. Treatment order (saline-OFF, cocaine-OFF, saline-ON, cocaine-ON) was counterbalanced between animals with a seven-day period between each treatment. Animals were first placed in the open field and connected to the stimulation for a 60 min period of habituation. They were then injected with either saline (0.9% NaCl, 1 ml/kg) or a low dose of cocaine (5 mg/kg, *s.c*.) and their locomotor activity was recorded for 60 minutes with DBS ON or OFF.

### Hot plate analgesia measurement

In another set of animals (*n* = 12), the effects of 2 h-STN DBS (at 0−, 30− or 130-Hz) for 5 consecutive days accordingly with the procedure used to test the effects of these frequencies on the compulsivity measures were tested on nociceptive responses after the first and the last sessions of STN DBS (Fig. S5).

In *n* = 6 animals that had completed the experiment for at least 2 months, basal nociceptive response was determined, regardless of their compulsivity status. Then, animals were subjected to 6 h-STN DBS (at 8-Hz) for 14 consecutive days accordingly with the procedure used to test the effects of this frequency applied during the escalation procedure. Their nociceptive responses were tested after the first and the last sessions of STN 8-Hz DBS (Fig. S10).

### Recording and analysis of STN LFP activity

LFP recordings were performed in similar operant chambers as above with no access to cocaine. They were further equipped with wires connected to the acquisition setup. Animals were connected to the interface and placed in the chamber where they could freely move. STN electric activity was recorded 15-min before and after extended cocaine access on days 1, 4, 8, 12 and 15 of the escalation protocol.

In another experiment (Fig. S8), LFPs were recorded (15-min) in naïve rats (n = 4) before and after the first 6-h session of 8-Hz STN DBS and after seven days of 8-Hz DBS. Signals were amplified and filtered using a Neuralynx8 amplifier. Data were acquired using Sciworks software (Datawave Tech, USA) with a sampling rate of 1kHz in the range of 1-475Hz. Signals were filtered off-line with a Chebyshev low pass filter (corner 98Hz, order 10, ripple 0.5) and a notch filter were applied, to remove 50Hz noise created by surrounding electrical devices, using Spike2 software (CED). Data were then carefully examined to ensure removal of electrical noise the specific activity of the STN (*i.e.* difference of potential between the two wires within the same STN) was then calculated and ultimately treated using Matlab (Mathworks) software. As such, the analysis was limited to the following frequency bands: 4-40 Hz and 65-90 Hz.

### Deep brain stimulation

DBS was delivered to the STN by a digital stimulator (DS8000, WPI) via a stimulus isolator (DLS100, WPI) and a rotating commutator (Plastics-One) wired to the implanted electrodes. Stimulation parameters were adapted from previous studies (*19*, *22*)). Briefly, individual stimulation intensity was determine using 130-Hz frequency and 80 μs pulse width stimulation. Intensity was progressively increased until the appearance of hyperkinetic movements. Stimulation intensity (50-150 μA) was setup just below the hyperkinetic movement threshold.

Before each behavioral session, animals were connected to the stimulation device, STN DBS was turned ON and stimulation intensity was progressively increased to reach the pre-determined stimulation parameters prior the start of the session. Depending on the condition tested, 8−, 30− or 130-Hz were applied during 1, 2 or 6 h.

### Histology

At the end of the experiment, the rats were euthanized with an *i*.*p*. injection of pentobarbital (Dolethal). Brains were removed and frozen into liquid isopentane (−80°C) to be further cut in 40 μm thick frontal slices with a cryostat. Histological controls for location of the electrodes were performed after staining with cresyl violet by an observer blind to treatment group (Figs. S2H, S4B, S5B, S7E, S8C, S9F, S10B). Animals with incorrect electrode placement (*n* = 11 in total, *n* = 3 for Fig. 2 and S2, 2 for Fig. 4 (70-Hz group), 3 for Fig. S5, 1 for S8 and 2 for Fig. S10), as well as those with catheter clogging (*n* = 4) were excluded from statistical analysis. The final *n* values are indicated in the figure legends and results section.

### Statistical analyses

Data are expressed as mean ± s.e.m. with the exact sample size indicated for each group in figure legends. Using Prism 6.0 (GraphPad) and Matlab (Mathworks) softwares, data were analyzed two-tailed *t* test, one- or two-way repeated measures ANOVAs, followed by Bonferroni *post hoc* test when applicable. Only *P*-values < 0.05 were considered significant.

## Supplementary Materials

Figure S1. Escalation of cocaine intake is a prerequisite to the onset of compulsive-like cocaine seeking behavior.

Figure S2. 130-Hz STN DBS has no long-lasting effects on compulsive-like cocaine seeking.

Figure S3. STN DBS does not affect seeking for natural reward during punished cocaine seeking.

Figure S4. 30-Hz STN DBS has no effect on rat’s locomotion.

Figure S5. STN DBS does not affect rat’s peripheral pain sensitivity.

Figure S6. Animals’ compulsivity status cannot be predicted before exposure to the punishment contingency.

Figure S7. ‘Shock-resistant’, but not ‘shock-sensitive’, rats display pathological STN low frequency oscillations during cocaine escalation, which are normalized by cocaine consumption.

Figure S8: 8-Hz STN DBS increases low frequency oscillations in naïve rats. Figure S9. STN DBS applied during escalation 2 in ‘shock-sensitive’ rats has no consequence on basal seeking and consummatory behaviors.

Figure S10. Repeated 8-Hz STN DBS does not affect rat’s peripheral pain sensitivity.

Table S1. Statistical analysis.

## General

The authors thank Drs. SH Ahmed, GF Koob, and D Robbe for critical reading of the manuscript and J Baurberg for technical support.

## Author contributions

M.D., C.B. and Y.P. designed and planned all experiments; M.D., A.T.C. and Y.P. performed experiments and analyzed data; M.D., C.B. and Y.P. wrote the paper. C.B. obtained the funding.

## Data and materials availability

Materials, datasets and protocols are available upon request to C.B.

