## supplementary figures for "Low frequency oscillatory activity of the subthalamic nucleus is a predictive biomarker of compulsive-like cocaine seeking"

### Supplementary Materials

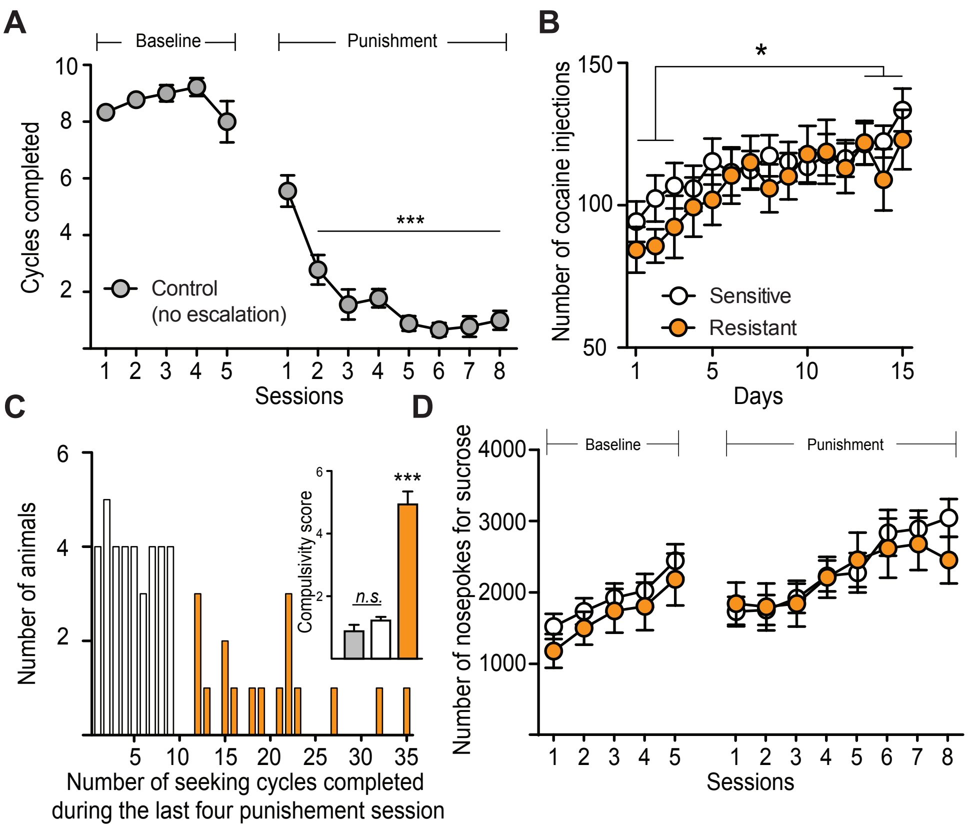


**Figure S1. Escalation of cocaine intake is a prerequisite to the onset of compulsive-like cocaine seeking behavior**. (**A**) Following training on the seeking-taking task, control animals (*n* = 9, not previously exposed to cocaine escalation) were subjected to five days of baseline cocaine seeking and then exposed to eight punishment sessions, during which foot shock (with no access to cocaine) was randomly delivered in 50% of the trials and cocaine for the other 50%. Punishment contingency completely suppressed cocaine seeking ($F_{(12,96)}$ = 90.66, $P < 0.0001$; Bonferroni post hoc: ***$P < 0.001$ baseline *vs.* punishment). (**B**) During the escalation protocol (6h/day for 15 days), future 'shock-sensitive' (white dots, *n* = 36) and 'shock-resistant' (orange dots, *n* = 17) rats escalated their cocaine intake in a similar manner (session effect: $F_{(14,714)}$ = 6.71, $P < 0.0001$; group effect: $F_{(1,51)}$ = 0.448, *n.s.*; Bonferroni post hoc: *$P < 0.05$). (**C**) Frequency distribution of the total number of cycles completed by all animals (those previously subjected to cocaine escalation) during the last four sessions of punishment, highlighting the bimodal distribution of individuals responding for cocaine under punishment. Inset: 'Shock-resistant' animals displayed a higher compulsivity score (*i.e.* mean number of cycles completed during the last four sessions of punishment) compared to 'shock-sensitive' and control rats ($F_{(2,59)}$ = 80.37, $P < 0.0001$; Bonferroni post hoc: ***$P < 0.001$). (**D**) Number of nosepokes for sucrose was similar for both groups during baseline (group effect: $F_{(1,51)}$ = 0.7797, *n.s.*) and punished cocaine seeking (group effect: $F_{(1,51)}$ = 0.0836, *n.s.*). Line and bar graphs indicate mean ± s.e.m.

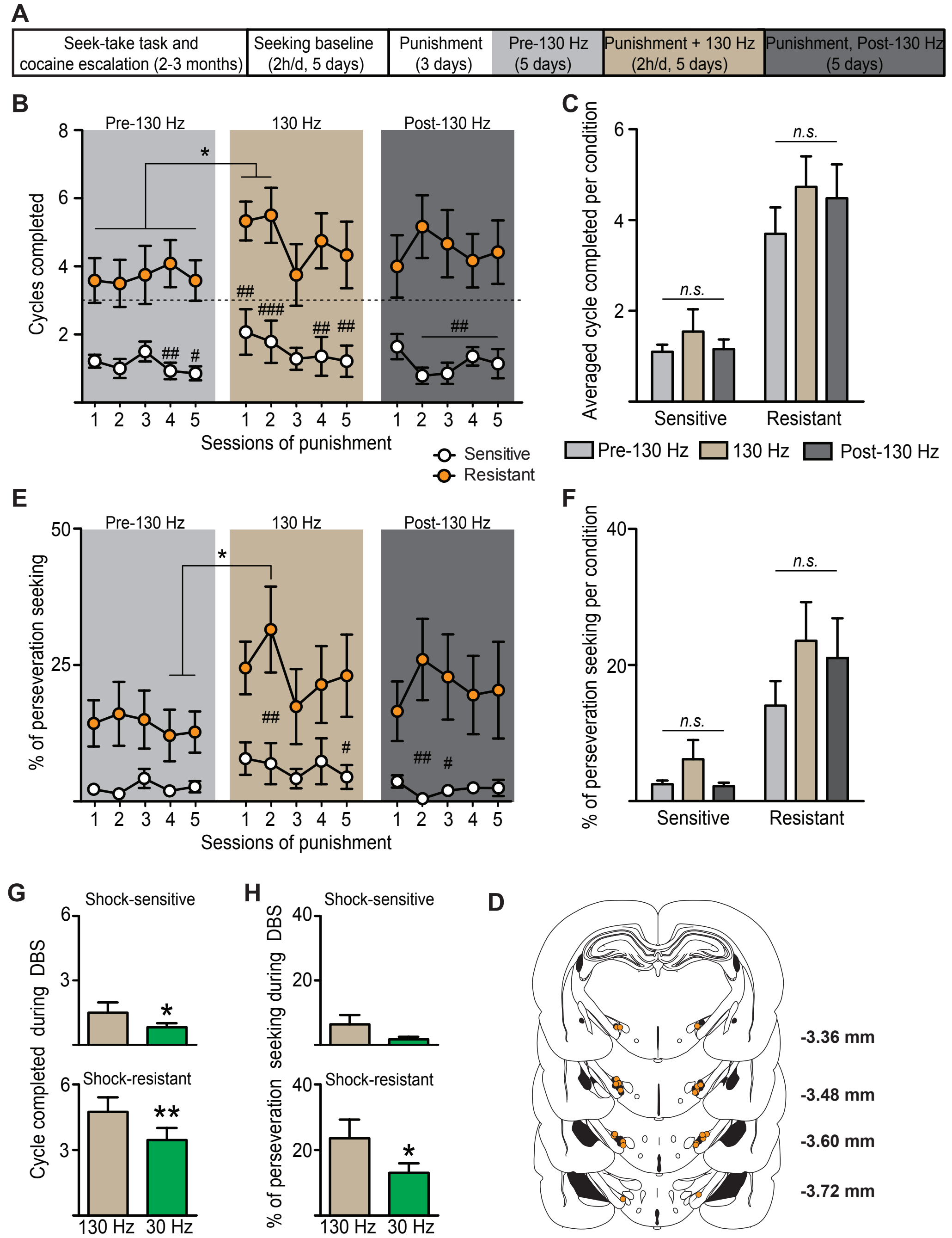


**Figure S2. 130-Hz STN DBS has no long-lasting effects on compulsive-like cocaine seeking.** (**A**) Experimental time course.After characterization of their compulsive status during the baseline-punishment sequence, animals were subjected to five days of 130-Hz STN DBS followed by 5 days with no DBS. (**B**) 130-Hz STN DBS acutely increased the number of seeking cycles completed by 'shock-resistant' rats (orange dots; $n$ = 12, DBS effect: $F_{(14,336)}$ = 1.764, $P$ = 0.0427), but had no effect in 'shock-sensitive' animals (white dots; $n$ = 14, group effect, $F_{(1,24)}$ = 26.01, $P$ < 0.0001; DBS x group effect: $F_{(14,336)}$ = 0.9112, *n.s.*; Bonferroni post hoc: *$P$ < 0.05; $^{\#}P$ < 0.05, $^{\#\#}P$ < 0.01, $^{\#\#\#}P$ < 0.001, resistant *vs*. sensitive). Dashed line indicates compulsivity threshold below which animals are considered shock-sensitive. (**C**) Averaged (5-session block) number of cycles completed by 'shock-sensitive' (left) and 'shock-resistant' (right) rats before (pale grey), during (brown) and after (dark grey) 130-Hz STN DBS (DBS effect: $F_{(2,48)} = 2.489$, *n.s.*; group effect: $F_{(1,24)} = 26.01$, $P < 0.0001$; DBS x group effect: $F_{(2,48)} = 0.6766$, *n.s.*). (**D**) 130-Hz STN DBS acutely increased the percentage of compulsive seeking lever presses performed by 'shock-resistant' rats (DBS effect: $F_{(14,336)} = 1.718$, $P = 0.05$), but had no effect in 'shock-sensitive' animals (group effect, $F_{(1,24)} = 15.84$, $P < 0.001$; DBS x group effect: $F_{(14,336)} = 0.9129$ *n.s.*; Bonferroni post hoc: $*P < 0.05$; $^{\#}P < 0.05$, $^{\#\#}P < 0.01$, resistant *vs*. sensitive). (**E**) Averaged (5-session block) percentage of compulsive seeking lever presses performed by ‘shock-sensitive’ (left) and ‘-resistant’ (right) rats before (pale grey), during (brown) and after (dark grey) 130-Hz STN DBS (group effect: $F_{(1,24)} = 15.84$, $P < 0.001$; DBS effect: $F_{(2,48)} = 2.973$, *n.s.*; DBS x group effect: $F_{(2,48)} = 1.019$, *n.s.*). (**F**) Averaged (5-session block) number of cycles completed by shock-sensitive (top) and -resistant (bottom) rats during 130-Hz and 30-Hz DBS ($t_{13} = 2.268$, $P < 0.05$ for ‘shock-sensitive’; $t_{11} = 3.578$, $P < 0.01$ for ‘shock-sensitive’; Bonferroni post hoc: $*P < 0.05$; $**P < 0.01$). (**G**) Averaged (5-session block) percentage of compulsive seeking lever presses performed by ‘shock-sensitive’ (top) and ‘shock-resistant’ (bottom) rats during 130-Hz and 30-Hz DBS ($t_{13} = 2.115$, *n.s.* for ‘shock-sensitive’; $t_{11} = 2.414$, $P < 0.05$ for ‘shock-sensitive’; Bonferroni post hoc: $*P < 0.05$). Line and bar graphs indicate mean ± s.e.m.

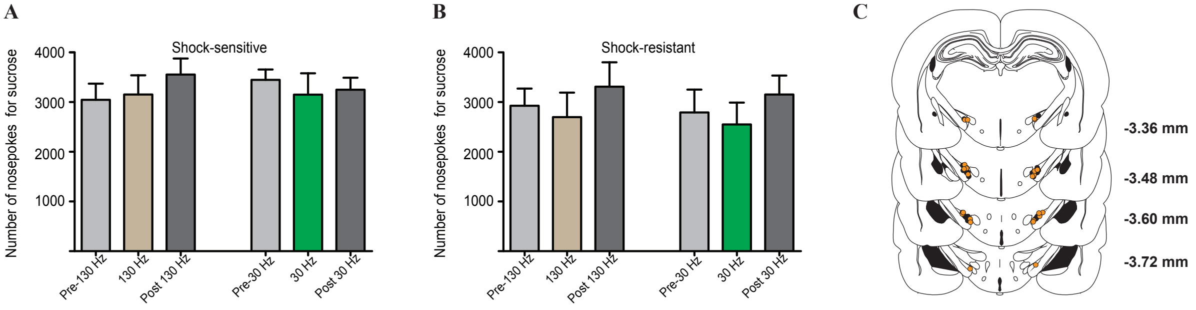


**Figure S3. STN DBS does not affect seeking for natural reward during punished cocaine seeking**. 30-Hz (green bars) or 130-Hz (brown bars) STN DBS during 5 punishment sessions had no effect on the number of nosepokes for sucrose in both ‘shock-sensitive’ (**A**, $n = 14$; $F_{(5,83)} = 0.573$, *n.s.*) and ‘shock-resistant’ animals (**B**, $n = 12$; $F_{(5,55)} = 2.099$, *n.s.*). Bar graphs indicate mean ± s.e.m. of 5 session-blocks. (**C**), Histologically verified electrode placements for 30-Hz and 130-Hz STN DBS also used in Fig. 2 (black dots: ‘shock-sensitive’, orange dots: ‘shock-resistant’). Numbers indicate distance from bregma (*52*).

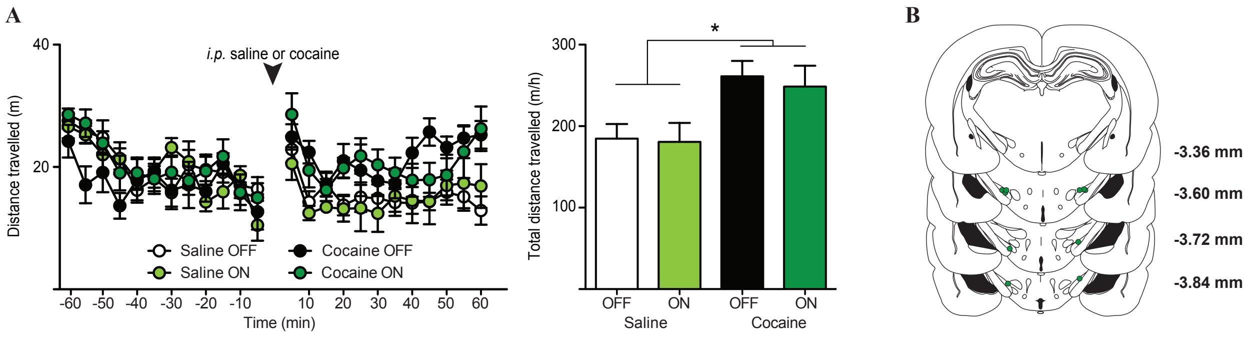


**Figure S4. 30-Hz STN DBS has no effect on rat's locomotion.** (**A**) Left: Time course of rats' (*n* = 6) locomotor activity (expressed as the distance travelled per bins of 5 minutes) during 60 minutes before and after an *i.p.* injection of saline (1 ml/kg, pale dots) or cocaine (5 mg/kg, dark dots). Right: When applied after an hour of habituation, 30-Hz STN DBS (green dots) affected neither rat's basal locomotion nor cocaine-induced hyperlocomotion ( $F_{(3,15)} = 10.19$, $P < 0.001$: Bonferroni post hoc: *$P < 0.05$). Line and bar graphs indicate mean ± s.e.m. (**B**) Histologically verified electrode placements (green dots). Numbers indicate distance from bregma (*52*).

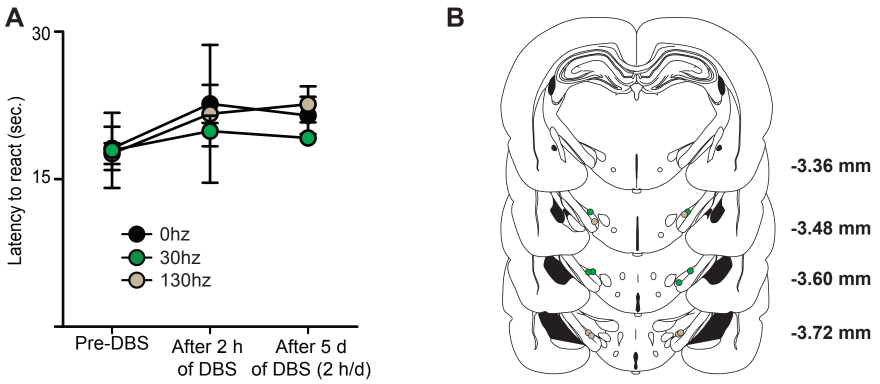


**Figure S5. STN DBS does not affect rat's peripheral pain sensitivity.** (**A**) Animals were subjected to 5 consecutive days of 2 h STN DBS (0 Hz, *n* = 3, black dots; 30-Hz, *n* = 3, green dots; 130-Hz, *n* = 3, brown dots). Nociceptive responses, measured as the latency to react on a hot plate via a video system, were evaluated before and after the first session of stimulation and after 5 daily sessions of stimulation. All groups displayed comparable latency to react (DBS effect: $F_{(2,12)} = 2.034$*, n.s.*; group effect: $F_{(2,6)} = 0.1722$*, n.s.*). Lines indicate mean ± s.e.m. (**B**) Histologically verified electrode placements for 30-Hz (green dots) and 130-Hz (brown dots) groups. Numbers indicate distance from bregma (*52*).

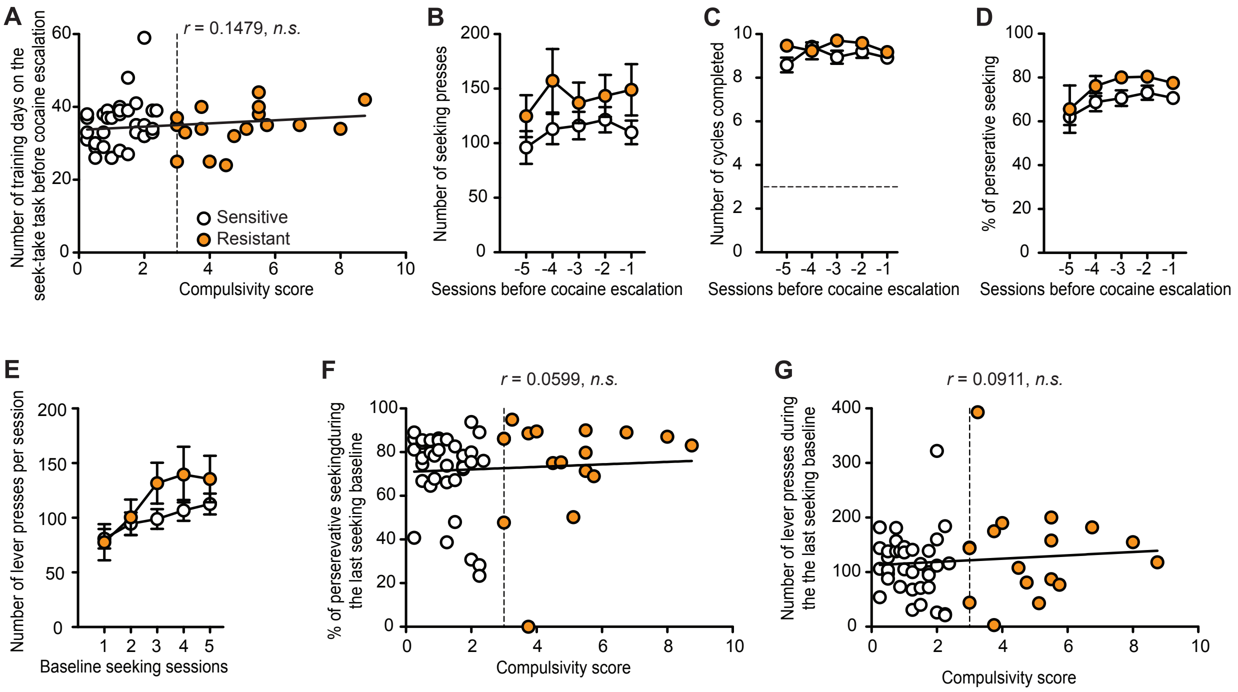


**Figure S6. Animals' compulsivity status cannot be predicted before exposure to the punishment contingency.** (**A**) No correlation was found between the number of training days on the seek-take task and the compulsivity score of 'shock-sensitive' (white dots) and 'shock-resistant' (orange dots) animals ($r$ = 0.1479, *n.s.*). (**B-D**) During the last five training sessions on the seek-take task (before being subjected to cocaine escalation protocol), both populations showed equivalent (**B**) level of lever presses (group effect: $F_{(1,51)}$ = 2.512*, n.s.*), (**C**) number of cycles completed (group effect: $F_{(1,51)}$ = 2.278*, n.s.*) and (**D**) percentage of compulsive seeking lever presses (group effect: $F_{(1,51)}$ = 2.844*, n.s.*). (**E**) Following cocaine escalation, 'shock-sensitive' and 'shock-resistant' rats displayed comparable number of lever presses during the five baseline seeking sessions (group effect: $F_{(1,51)}$ = 1.241*, n.s.*). (**F-G**) No correlation was found between the percentage of compulsive seeking lever presses (**F**: $r$ = 0.0599, *n.s.*) or the number of lever presses (**G**: $r$ = 0.0911, *n.s.*) performed during the last baseline seeking session and animals' compulsivity score. Lines indicate mean ± s.e.m.

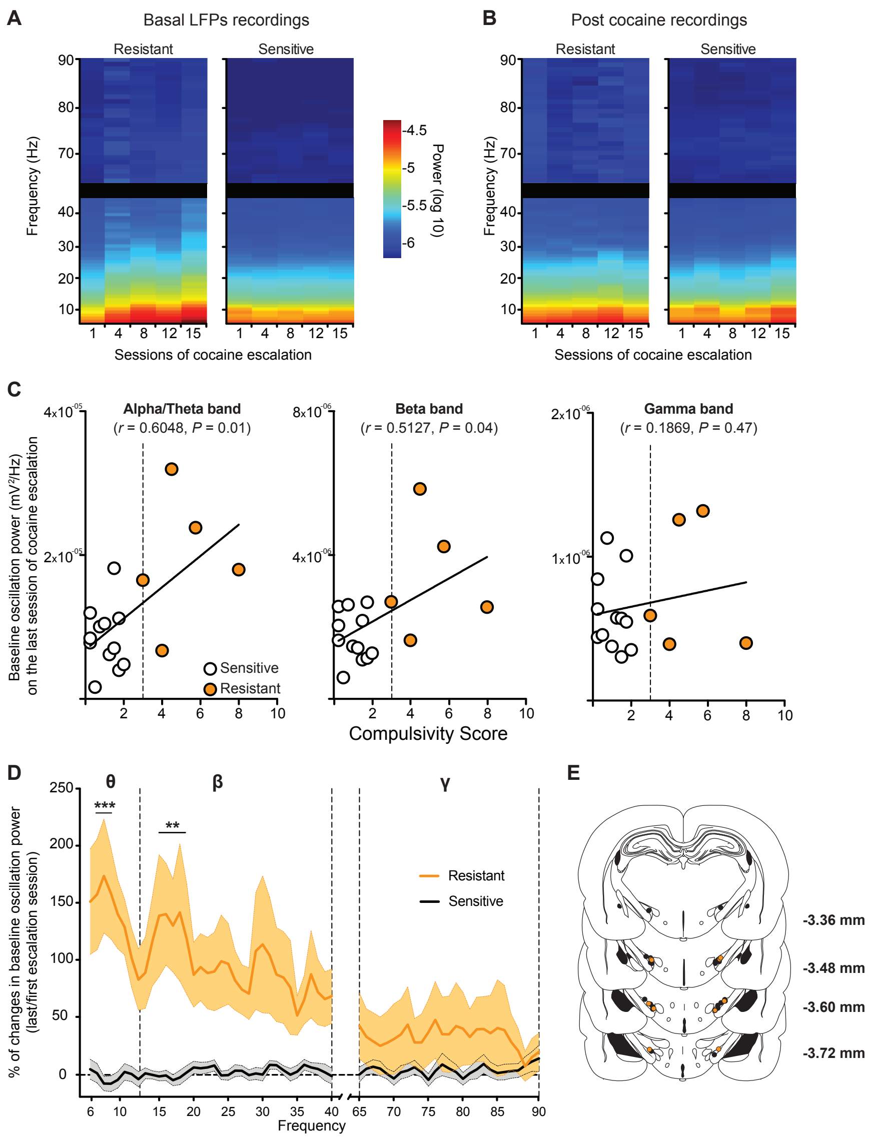


**Figure S7. ‘Shock-resistant’, but not ‘shock-sensitive’, rats display pathological STN low frequency oscillations during cocaine escalation, which are normalized by cocaine consumption.** (**A-B**) Session-frequency raw power spectrums showing basal (**A**, before cocaine) and after 6 h of cocaine intake (**B**, post cocaine) LFPs power in ‘shock-resistant’ (left panels, *n* = 5) and ‘shock-sensitive’ (right panels, *n* = 12) animals. Values indicate logarithmic quantification of the power spectrum. (**C**) Animals’ compulsivity score was correlated with the baseline STN activity recorded before the last escalation session in the alpha/theta band (left panel: $r = 0.6049$, $P = 0.01$) and the beta band (middle panel: $r = 0.5127$, $P = 0.04$) but not in the gamma band (right panel: $r = 0.1869$, *n.s.*). (**D**) Mean percentage of STN basal oscillatory power changes between the last and the first cocaine escalation sessions for both populations. While cocaine escalation has no effect on oscillatory power for ‘shock-sensitive’ animals (session x frequency effect: $F_{(62,693)}$ =

1.091, *n.s.*, black line), it increases low frequency oscillations in 'shock-resistant' rats (orange line: session x frequency effect: $F_{(62,252)} = 1.548$, $P = 0.01$, Bonferroni post hoc: $**P < 0.01$, $***P < 0.001$ *vs*. first cocaine escalation session) notably in the alpha/theta (7-9 Hz) and low beta bands (15-19 Hz). Light-shaded areas indicate s.e.m. (**D**) Histologically verified electrode placements for LFPs recordings also used in Fig. 3 (black dots: 'shock-sensitive', orange dots: 'shock-resistant'). Numbers indicate distance from bregma (*52*).

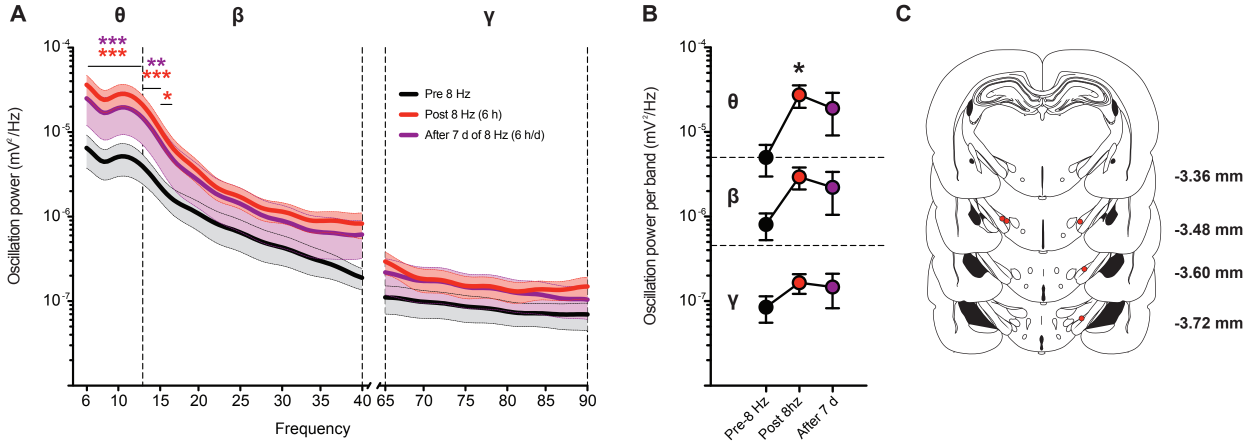


**Figure S8: 8-Hz STN DBS increases low frequency oscillations in naïve rats.** (**A**) Power spectrum of STN LFPs recorded before (black line) and after (red line) the first 6h session of 8-Hz STN DBS and after seven days of 6h 8-Hz STN DBS session (purple line). Acute and chronic 8-Hz DBS increases low, but not high, frequency oscillations ($n$ = 5, session x frequency effect: $F_{(276,1112)} = 3.461$, $P < 0.0001$, Bonferroni post hoc: *$P < 0.05$, **$P < 0.01$, ***$P < 0.001$ *vs*. pre-8-Hz, color-coded) notably in the alpha/theta (6-13 Hz) and low beta bands (14-16 Hz). Light-shaded areas indicate s.e.m. (**B**) Band-specific quantifications of LFPs power before and after one or seven sessions of 6h 8-Hz STN DBS (alpha/theta: $F_{(2,8)} = 4.465$, $P < 0.05$, beta: $F_{(2,8)} = 2.846$, *n.s.*; gamma: $F_{(2,8)} = 1.201$, *n.s.*). (**C**) Histologically verified electrode placements. Numbers indicate distance from bregma (*52*).

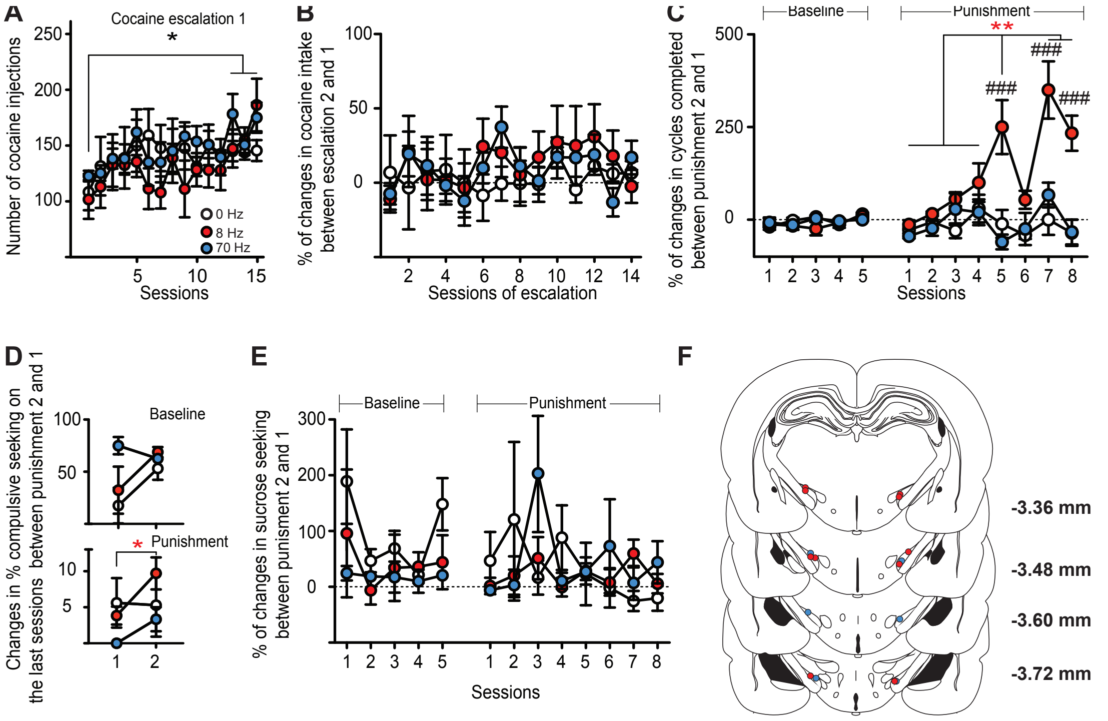


**Figure S9. STN DBS applied during escalation 2 in 'shock-sensitive' rats has no consequence on basal seeking and consummatory behaviors.** (**A**) All groups (0 Hz, $n$ = 5, white dots; 8-Hz, $n$ = 5, red dots; 70 Hz, $n$ = 3, blue dots) displayed comparable cocaine intake during escalation 1 (session effect: $F_{(14,140)} = 3.136$, $P < 0.001$; group effect: $F_{(2,10)} = 0.3937$, *n.s.*; Bonferroni post hoc: $*P < 0.05$). (**B**) STN DBS applied during cocaine escalation 2 did not affect the dynamic of intake observed during the cocaine escalation 1, since the percentage of change between escalation 1 and 2 is not significantly different in all groups (session effect: $F_{(13,140)} = 0.601$, *n.s.*; group effect: $F_{(2,140)} = 1.079$, *n.s.*). (**C**) STN DBS applied during cocaine escalation 2 did not affect the dynamic of basal cocaine seeking observed during the baseline 2, since the percentage of change between baseline 1 and 2 is not significantly different in all groups (session x group effect: $F_{(8,40)} = 1.532$, *n.s*). In contrast, only 8-Hz STN DBS changed the dynamic of compulsive-like cocaine seeking during punished sessions (session effect: $F_{(7,70)} = 9.131$, $P < 0.0001$; group effect: $F_{(2,10)} = 9.532$, $P < 0.01$; Bonferroni post hoc: $**P < 0.01$ for 8-Hz; $^{\#\#}P < 0.01$, $^{\#\#\#}P < 0.001$ 8-Hz *vs*. control). (**D**) All groups exhibited comparable level of compulsive seeking lever presses during the last session of baseline seeking before and after STN DBS applied during cocaine escalation 2 (session effect: $F_{(2,10)} = 1.791$, *n.s.*). In contrast, 8-Hz stimulated animals performed more compulsive seeking lever presses during the last punished seeking session $F_{(2,10)} = 6.342$, $P < 0.05$, Bonferroni post hoc: $*P < 0.05$ for 8-Hz). (**E**) STN DBS applied during cocaine escalation 2 did not affect the dynamic of sucrose seeking observed during punishment 2, since the percentage of change between punishment 1 and 2 is not significantly different in all groups (session effect: $F_{(12,130)}$ =

0.8742, *n.s.*; group effect: $F_{(2,130)} = 0.9482$, *n.s.*). Line and bar graphs indicate mean ± s.e.m. (**F**) Histologically verified electrode placements for 8-Hz (red dots) and 70 Hz (blue dots) STN DBS also used in Fig. 4. Numbers indicate distance from bregma (*52*).

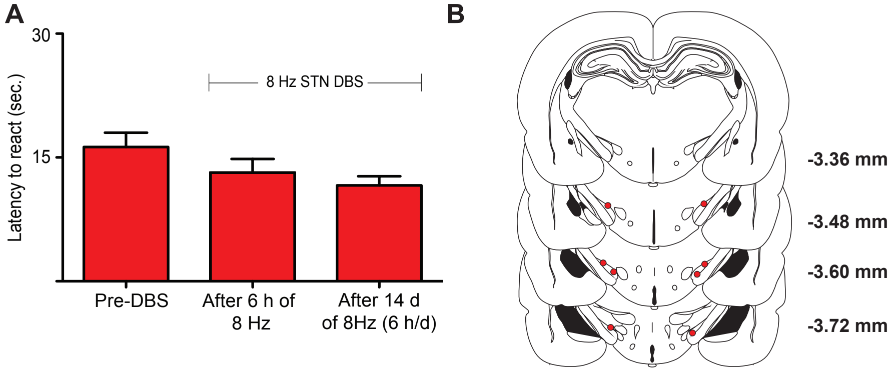


**Figure S10. Repeated 8-Hz STN DBS does not affect rat's peripheral pain sensitivity.** (**A**) Animals were subjected to 14 consecutive days of 6 h of 8-Hz STN DBS. Nociceptive responses, measured as the latency to react on a hot plate via a video system, were evaluated before and after the first session of stimulation and after 14 daily sessions of stimulation. Latency to react to a hot plate was not affected by 14 consecutives days of 6 h of 8-Hz STN DBS ($n$ = 4, $F_{(2,6)}$ = 3.34, *n.s.*). Bar graphs indicate mean ± s.e.m. (**B**) Histologically verified electrode placements (yellow dots). Numbers indicate distance from bregma (*52*).

### Table S1: Statistical analyses

| Figure | Conditions<br>(n = number of animals per condition) | Statistical analyses |
|---|---|---|
| **1B** | Sensitive (n =36)<br><br>Resistant (n = 17) | 2-way repeated measure ANOVA:<br>* Sessions: $F_{12,612} = 117.2$, $P < 0.0001$.<br>* Group: $F_{1,51} = 30.12$, $P < 0.0001$.<br>* Sessions x Group: $F_{12,612} = 15.85$, $P < 0.0001$.<br><br>2-way repeated measure ANOVA on baseline sessions:<br>* Sessions: $F_{4,204} = 8.831$, $P < 0.0001$.<br>* Group: $F_{1,51} = 0.559$, $P = 0.4582$.<br>* Sessions x Group: $F_{4,204} = 2.704$, $P = 0.0316$.<br><br>2-way repeated measure ANOVA for punishment sessions:<br>* Sessions: $F_{7,357} = 23.76$, $P < 0.0001$.<br>* Group: $F_{1,51} = 69.91$, $P < 0.0001$.<br>* Sessions x Group: $F_{7,357} = 7.927$, $P < 0.0001$.<br><br>Bonferroni *post-hoc* test:<br>* Sensitive vs. Resistant:<br>$P < 0.01$ for session 2 (power = 0.84).<br>$P < 0.001$ for sessions 3-8 ($0.92 \leq$ power $\leq 1$). |
| **1C** | | 2-way repeated measure ANOVA:<br>* Sessions: $F_{1,51}$= 355.9, $P < 0.0001$.<br>* Group: $F_{1,51} = 32.62$, $P < 0.0001$.<br>* Sessions x Group: $F_{1,51} = 35.63$, $P < 0.0001$.<br><br>Bonferroni *post-hoc* test:<br>* Baseline vs. Punishment:<br>$P < 0.001$ for Sensitive and Resistant (power = 1).<br>* Sensitive vs. Resistant:<br>$P < 0.001$ for punishment (power = 1). |
| **1D** | | 2-way repeated measure ANOVA:<br>* Sessions: $F_{12,612} = 122.7$, $P < 0.0001$.<br>* Group: $F_{1,51} = 11.85$, $P = 0.0012$.<br>* Sessions x Group: $F_{12,612} = 4.605$, $P < 0.0001$.<br><br>2-way repeated measure ANOVA on baseline sessions:<br>* Sessions: $F_{4,204} = 12.36$, $P < 0.0001$.<br>* Group: $F_{1,51} = 0.009$, $P = 0.9225$.<br>* Sessions x Group: $F_{4,204} = 2.169$, $P = 0.0737$.<br><br>2-way repeated measure ANOVA for punishment sessions:<br>* Sessions: $F_{7,357} = 31.71$, $P < 0.0001$. |

| | | |
|---|---|---|
| | Sensitive (n =36)<br><br>Resistant (n = 17) | * Group: $F_{1,51}$ = 33.79, $P$ < 0.0001.<br>* Sessions x Group: $F_{7,357}$ = 1.686, $P$ = 0.1110.<br><br>Bonferroni *post-hoc* test:<br>* Sensitive vs. Resistant:<br>$P$ < 0.01 for session 2-3, 6, 8 (0.71 ≤ power ≤ 0.97).<br>$P$ < 0.001 for sessions 4-5, 7 (0.98 ≤ power ≤ 1). |
| **1E** | | 2-way repeated measure ANOVA:<br>* Sessions: $F_{1,51}$= 327.3, $P$ < 0.0001.<br>* Group: $F_{1,51}$ = 8.323, $P$ = 0.0057.<br>* Sessions x Group: $F_{1,51}$ = 4.184, $P$ = 0.046.<br><br>Bonferroni *post-hoc* test:<br>* Baseline vs. Punishment:<br>$P$ < 0.001 for Sensitive and Resistant (power = 1).<br>* Sensitive vs. Resistant:<br>$P$ = 0.0014 for punishment (power = 1). |
| **2B** | Sensitive (n =14)<br><br>Resistant (n = 12) | 2-way repeated measure ANOVA:<br>* DBS: $F_{14,336}$ = 3.494, $P$ < 0.0001.<br>* Group: $F_{1,24}$ = 33.67, $P$ < 0.0001.<br>* DBS x Group: $F_{14,336}$ = 2.848, $P$ = 0.0005.<br><br>2-way repeated measure ANOVA on baseline sessions:<br>* DBS: $F_{4,96}$ = 1,997, $P$ = 0.1011.<br>* Group: $F_{1,24}$ = 40.38, $P$ < 0.0001.<br>* DBS x Group: $F_{4,96}$ = 2.085, $P$ = 0.0887.<br><br>2-way repeated measure ANOVA on ON sessions:<br>* DBS: $F_{4,96}$ = 1.626, $P$ = 0.1740.<br>* Group: $F_{1,24}$ = 22.13, $P$ < 0.0001.<br>* DBS x Group: $F_{4,96}$ = 2.312, $P$ = 0.0631.<br><br>2-way repeated measure ANOVA on OFF sessions:<br>* DBS: $F_{4,96}$ = 0.5915, $P$ = 0.6696.<br>* Group: $F_{1,24}$ = 17.49, $P$ = 0.0003.<br>* DBS x Group: $F_{4,96}$ = 0.9552, $P$ = 0.4358.<br><br>Bonferroni *post-hoc* test:<br>* Sensitive vs. Resistant:<br>$P$ < 0.05 for ON 2,4 (0.94 ≤ power ≤ 0.95).<br>OFF 3 (power = 0.91).<br>$P$ < 0. 01 for OFF 1,4 (0.92 ≤ power ≤ 0.98).<br>$P$ < 0. 001 for baseline 1-5 (0.95 ≤ power ≤ 1).<br>ON 1,3 (power = 0.99). |

| | | |
|---|---|---|
| | | * Resistant:<br>$P < 0.05$ for ON 1 vs. Baseline 2 (power = 1).<br>OFF 2 (power = 1).<br>ON 2 vs. Baseline 1,5 (power = 1).<br>ON 3 vs. Baseline 2 (power = 1).<br>ON 4 vs. Baseline 1,3 (power = 1).<br>ON 5 vs. OFF 4 (power = 0,98).<br>$P < 0.01$ for ON 1 vs. ON 5 (power= 1).<br>ON 2 vs. Baseline 4 (power = 1).<br>ON 3 vs. ON 5 (power = 0,99).<br>ON 4 vs. Baseline 5 (power = 1).<br>ON 5 vs. ON 1,3 (power = 1).<br>OFF 1 (power = 1).<br>$P < 0.001$ for ON 2 vs. Baseline 2 (power = 1).<br>ON 4 vs. Baseline 2,4 (power = 1).<br>ON 5 vs. Baseline 1-5 (power = 1). |
| **2C** | Sensitive (n =14)<br><br>Resistant (n = 12) | 2-way repeated measure ANOVA:<br>* DBS: $F_{2,48} = 9.416$, $P = 0.0004$.<br>* Group: $F_{1,24} = 33.67$, $P < 0.0001$.<br>* DBS x Group: $F_{2,48} = 5.75$, $P = 0.0058$.<br><br>Bonferroni *post-hoc* test:<br>* Sensitive vs. Resistant:<br>$P < 0.001$ for Baseline, ON, OFF (power = 0.99).<br>* Resistant:<br>$P < 0.001$ for Baseline vs. ON, OFF (power = 1). |
| **2D** | | 2-way repeated measure ANOVA:<br>* DBS: $F_{14,336} = 2.6$, $P = 0.0014$.<br>* Group: $F_{1,24} = 22.32$, $P < 0.0001$.<br>* DBS x Group: $F_{14,336} = 2.88$, $P = 0.0004$.<br><br>2-way repeated measure ANOVA on baseline sessions:<br>* DBS: $F_{4,96} = 0.1523$, $P = 0.9616$.<br>* Group: $F_{1,24} = 20.08$, $P = 0.0002$.<br>* DBS x Group: $F_{4,96} = 0.8964$, $P = 0.4693$.<br><br>2-way repeated measure ANOVA on ON sessions:<br>* DBS: $F_{4,96} = 3.792$, $P = 0.0019$.<br>* Group: $F_{1,24} = 16.67$, $P = 0.0004$.<br>* DBS x Group: $F_{4,96} = 4.607$, $P = 0.0019$.<br><br>2-way repeated measure ANOVA on OFF sessions:<br>* DBS: $F_{4,96} = 0.6436$, $P = 0.6328$.<br>* Group: $F_{1,24} = 11.95$, $P = 0.0021$.<br>* DBS x Group: $F_{4,96} = 0.9545$, $P = 0.4362$. |

| | | |
|---|---|---|
| | Sensitive (n =14)<br><br>Resistant (n = 12) | Bonferroni *post-hoc* test:<br>* Sensitive vs. Resistant:<br>$P < 0.05$ for Baseline 1 (power = 0.89).<br>$P < 0.01$ for Baseline 3 (power = 0.96).<br>ON 3 (power = 0.93).<br>$P < 0.001$ for Baseline 2,4-5 ($0.96 \leq$ power $\leq 1$).<br>ON 1 (power = 1).<br><br>* Resistant:<br>$P < 0.05$ for ON 1 vs. ON 2 (power = 1).<br>OFF 2 (power = 1).<br>ON 3 vs. ON 4 (power = 0.97).<br>ON 4 vs. Baseline 1,3 ($0.99 \leq$ power $\leq 1$).<br>ON 5 vs. Baseline 1,3 ($0.99 \leq$ power $\leq 1$).<br>$P < 0.01$ for ON 1 vs. ON 4-5 (power = 1).<br>ON 2 vs. Baseline 2,4 ($0.99 \leq$ power $\leq 1$).<br>ON 4 vs. Baseline 5 (power = 0.98).<br>ON 5 vs. ON 1,3 ($0.94 \leq$ power $\leq 1$).<br>OFF 1 (power = 1).<br>$P < 0.001$ for ON 4 vs. Baseline 2,4 (power = 1).<br>ON 5 vs. Baseline 2,4-5 ($0.97 \leq$ power $\leq 1$). |
| **2E** | | 2-way repeated measure ANOVA:<br>* DBS: $F_{2,48} = 6.168$, $P = 0.0041$.<br>* Group: $F_{1,24} = 22.32$, $P < 0.0001$.<br>* DBS x Group: $F_{2,48} = 5.389$, $P = 0.0077$.<br><br>Bonferroni *post-hoc* test:<br>* Sensitive vs. Resistant:<br>$P < 0.01$ for ON, OFF ($0.98 \leq$ power $\leq 0.99$).<br>$P < 0.001$ for Baseline (power = 1).<br>* Resistant:<br>$P < 0.01$ for Baseline vs. ON, OFF (power = 1). |
| **3A** | Sensitive (n =12)<br><br>Resistant (n = 5) | 2-way repeated measure ANOVA on escalation sessions:<br>* Sessions: $F_{14,210} = 6.675$, $P < 0.0001$.<br>* Group: $F_{1,15} = 0.813$, $P = 0.3815$.<br>* Sessions x Group: $F_{14,210} = 0.7089$, $P = 0.7643$.<br><br>Bonferroni *post-hoc* test:<br>$P < 0.05$ for sessions 1-4 vs. sessions 12-15. |
| **3B** | | 2-way repeated measure ANOVA:<br>* Sessions: $F_{12,180} = 66.21$, $P < 0.0001$.<br>* Group: $F_{1,15} = 20.81$, $P = 0.0004$.<br>* Sessions x Group: $F_{12,180} = 7.305$, $P < 0.0001$. |

| | | |
|---|---|---|
| | | 2-way repeated measure ANOVA for baseline sessions:<br>* Sessions: $F_{4,60}$ = 2.191, $P$ = 0.0807.<br>* Group: $F_{1,15}$ = 0.2039, $P$ = 0.658.<br>* Sessions x Group: $F_{4,60}$ = 1.067, $P$ = 0.3809.<br><br>2-way repeated measure ANOVA for punishment sessions:<br>* Sessions: $F_{7,105}$ = 13.91, $P$ < 0.0001.<br>* Group: $F_{1,15}$ = 25.32, $P$ = 0.0001.<br>* Sessions x Group: $F_{7,105}$ = 3.947, $P$ = 0.0007.<br><br>Bonferroni *post-hoc* test:<br>* Sensitive vs. Resistant:<br>$P$ < 0.05 for session 4 (power = 0.65).<br>$P$ < 0.01 for session 6 (power = 0.73).<br>$P$ < 0.001 for session 5,7,8 (0.83 ≤ power ≤ 1). |
| **3D** | Sensitive (n =12)<br><br>Resistant (n = 5) | 2-way repeated measure ANOVA:<br>* Sessions: $F_{9,135}$ = 1.981, $P$ = 0.0461.<br>* Group: $F_{1,15}$ = 1.841, $P$ = 0.1949.<br>* Sessions x Group: $F_{9,135}$ = 4.059, $P$ = 0.0001.<br><br>2-way repeated measure ANOVA for baseline recordings:<br>* Sessions: $F_{4,60}$ = 5.481, $P$ = 0.0008.<br>* Group: $F_{1,15}$ = 5.454, $P$ = 0.0338.<br>* Sessions x Group: $F_{4,60}$ = 7.651, $P$ < 0.0001.<br><br>2-way repeated measure ANOVA for post-cocaine recordings:<br>* Sessions: $F_{4,60}$ = 0.4829, $P$ = 0.7482.<br>* Group: $F_{1,15}$ = 0.1205, $P$ = 0.7333.<br>* Sessions x Group: $F_{4,60}$ = 1.021, $P$ = 0.4036.<br><br>Bonferroni *post-hoc* test:<br>* Resistant:<br>$P$ < 0.05 for session 1 vs. session 4 (power = 0.98).<br>$P$ < 0.01 for session 1 vs. session 12 (power = 0.59).<br>session 4 vs. session 15 (power = 0.68).<br>$P$ < 0.001 for session 1 vs. session 8 (power = 0.72).<br>vs. session 15 (power = 0.91).<br>* Before vs. After:<br>$P$ < 0.001 for session 15 for resistant (power = 0.8).<br><br>* Sensitive vs. Resistant:<br>$P$ < 0.01 for session 15 (power = 0.75). |
| **3E** | | 2-way repeated measure ANOVA:<br>* Sessions: $F_{9,135}$ = 2.456, $P$ = 0.0127.<br>* Group: $F_{1,15}$ = 3.387, $P$ = 0.0856. |

| | | |
|---|---|---|
| | Sensitive (n =12) | * Sessions x Group: $F_{9,135}$ = 3.514, $P$ = 0.0006.<br><br>2-way repeated measure ANOVA for baseline recordings:<br>* Sessions: $F_{4,60}$ = 4.119, $P$ = 0.0051.<br>* Group: $F_{1,15}$ = 6.035, $P$ = 0.0267.<br>* Sessions x Group: $F_{4,60}$ = 4.427, $P$ = 0.0033.<br><br>2-way repeated measure ANOVA for post-cocaine recordings:<br>* Sessions: $F_{4,60}$ = 0.2894, $P$ = 0.8838.<br>* Group: $F_{1,15}$ = 0.9516, $P$ = 0.3448.<br>* Sessions x Group: $F_{4,60}$ = 0.6136, $P$ = 0.6545.<br><br>Bonferroni *post-hoc* test:<br>* Resistant:<br>$P$ < 0.05 for session 1 vs. sessions 12 (power = 0.83).<br>$P$ < 0.01 for session 1 vs. sessions 4 (power = 0.81).<br>vs. sessions 8 (power = 0.94).<br>$P$ < 0.001 for session 1 vs. session 15 (power = 0.88).<br>* Before vs. After:<br>$P$ < 0. 001 for session 15 for resistant (power = 0.58).<br>* Sensitive vs. Resistant:<br>$P$ < 0.01 for session 15 (power = 0.71). |
| **3F** | Resistant (n = 5) | 2-way repeated measure ANOVA:<br>* Sessions: $F_{9,135}$ = 1.533, $P$ = 0.1424.<br>* Group: $F_{1,15}$ = 1.348, $P$ = 0.2638.<br>* Sessions x Group: $F_{9,135}$ = 1.393, $P$ = 0.1972.<br><br>2-way repeated measure ANOVA for baseline recordings:<br>* Sessions: $F_{4,60}$ = 0.9093, $P$ = 0.4644.<br>* Group: $F_{1,15}$ = 1.929, $P$ = 0.1851.<br>* Sessions x Group: $F_{4,60}$ = 1.393, $P$ = 0.2472.<br><br>2-way repeated measure ANOVA for post-cocaine recordings:<br>* Sessions: $F_{4,60}$ = 2.627, $P$ = 0.0432.<br>* Group: $F_{1,15}$ = 0.8765, $P$ = 0.364.<br>* Sessions x Group: $F_{4,60}$ = 1.706, $P$ = 0.1605. |
| **4B** | 0 Hz (n =5)<br><br>8-Hz (n = 5)<br><br>70 Hz (n =3) | 2-way repeated measure ANOVA:<br>* Sessions: $F_{12,120}$ = 89.94, $P$ < 0.0001.<br>* Group: $F_{2,10}$ = 0.168, $P$ = 0.8479.<br>* Sessions x Group: $F_{24,120}$ = 1.091, $P$ = 0.3644.<br><br>2-way repeated measure ANOVA for baseline sessions:<br>* Sessions: $F_{4,40}$ = 2.185, $P$ = 0.0881.<br>* Group: $F_{2,10}$ = 0.449, $P$ = 0.6505.<br>* Sessions x Group: $F_{8,40}$ = 1.209, $P$ = 0.3187. |

| | | |
|---|---|---|
| | | 2-way repeated measure ANOVA for punishment sessions:<br>* Sessions: $F_{7,70} = 29.61$, $P < 0.0001$.<br>* Group: $F_{2,10} = 0.2029$, $P = 0.8196$.<br>* Sessions x Group: $F_{14,70} = 1.279$, $P = 0.2423$. |
| **4C** | | 2-way repeated measure ANOVA on escalation sessions:<br>* Sessions: $F_{13,130} = 5.302$, $P < 0.0001$.<br>* Group: $F_{2,10} = 2.142$, $P = 0.1681$.<br>* Sessions x Group: $F_{26,130} = 0.639$, $P = 0.9079$.<br><br>Bonferroni *post-hoc* test:<br>$P < 0.05$ for session 1 vs. sessions 9,12.<br>2 vs. sessions 9-10, 13-14.<br>$P < 0.001$ for session 1 vs. sessions 10-14. |
| **4D** | 0 Hz (n =5)<br><br>8-Hz (n = 5)<br><br>70 Hz (n =3) | 2-way repeated measure ANOVA:<br>* Sessions: $F_{12,120} = 40.94$, $P < 0.0001$.<br>* Group: $F_{2,10} = 4.16$, $P = 0.0485$.<br>* Sessions x Group: $F_{24,120} = 3.168$, $P < 0.0001$.<br><br>2-way repeated measure ANOVA for baseline sessions:<br>* Sessions: $F_{4,40} = 3.52$, $P = 0.0149$.<br>* Group: $F_{2,10} = 0.529$, $P = 0.6047$.<br>* Sessions x Group: $F_{8,40} = 1.055$, $P = 0.4129$.<br><br>2-way repeated measure ANOVA for punishment sessions:<br>* Sessions: $F_{7,70} = 9.108$, $P < 0.0001$.<br>* Group: $F_{2,10} = 10.48$, $P = 0.0035$.<br>* Sessions x Group: $F_{14,70} = 1.715$, $P = 0.072$.<br><br>Bonferroni *post-hoc* test:<br>* 0 Hz vs. 8-Hz:<br>$P < 0.05$ for sessions 4,6,8 ($0.7 \leq$ power $\leq 0.99$).<br>* 70 Hz vs. 8-Hz:<br>$P < 0.05$ for sessions 4, 8 ($0.83 \leq$ power $\leq 0.96$).<br>$P < 0.01$ for session 5 (power = 0.78). |
| **4E** | | 2-way repeated measure ANOVA:<br>* Sessions: $F_{1,10} = 5.4$, $P = 0.0425$.<br>* Group: $F_{2,10} = 11.22$, $P = 0.0028$.<br>* Sessions x Group: $F_{2,10} = 7.705$, $P = 0.0094$.<br><br>Bonferroni *post-hoc* test:<br>* 8-Hz (after):<br>$P < 0.001$ vs. 0 Hz (power = 0.91).<br>$P < 0.001$ vs. 70 Hz (power = 0.99).<br>* Before vs. After: |

| | | |
|---|---|---|
| | | $P < 0.01$ for 8-Hz (power = 0.99). |
| **S1A** | | 1-way repeated measure ANOVA:<br>$F_{12,96} = 90.66$, $P < 0.0001$.<br><br>Bonferroni *post-hoc* test:<br>$P < 0.001$ for baseline sessions 1-5 vs. punishment sessions 2-8. |
| **S1B** | | 2-way repeated measure ANOVA on escalation sessions:<br>* Sessions: $F_{14,714} = 6.71$, $P < 0.0001$.<br>* Group: $F_{1,51} = 0.4484$, $P = 0.5061$.<br>* Sessions x Group: $F_{14,714} = 0.71$, $P = 0.7262$.<br><br>Bonferroni *post-hoc* test:<br>$P < 0.01$ for sessions 1-3 vs. sessions 13-15. |
| **S1C** | Control (n = 9)<br><br>Sensitive (n = 36)<br><br>Resistant (n = 17) | Kolmogorov-Smirnov test:<br>Control vs. Sensitive: $D = 0.3056$, $P = 0.5121$.<br>Control vs. Resistant: $D = 1$, $P \leq 0.0001$.<br>Sensitive vs. Resistant: $D = 1$, $P \leq 0.0001$.<br><br>1-way ANOVA:<br>$F_{2,59} = 80.37$, $P < 0.0001$.<br><br>Bonferroni *post-hoc* test:<br>$P < 0.001$ Resistant vs. Control, Sensitive. |
| **S1D** | | 2-way repeated measure ANOVA:<br>* Sessions: $F_{12,612} = 9.633$, $P < 0.0001$.<br>* Group: $F_{1,51} = 0.291$, $P = 0.5919$.<br>* Sessions x Group: $F_{12,612} = 0.473$, $P = 0.9306$.<br><br>2-way repeated measure ANOVA for baseline sessions:<br>* Sessions: $F_{4,204} = 7.561$, $P < 0.0001$.<br>* Group: $F_{1,51} = 0.7797$, $P = 0.3814$.<br>* Sessions x Group: $F_{4,204} = 0.0533$, $P = 0.9947$.<br><br>2-way repeated measure ANOVA for punishment sessions:<br>* Sessions: $F_{7,357} = 9.399$, $P < 0.0001$.<br>* Group: $F_{1,51} = 0.08357$, $P = 0.7737$.<br>* Sessions x Group: $F_{7,357} = 0.7343$, $P = 0.643$. |
| **S2B** | Sensitive (n =14)<br><br>Resistant (n = 12) | 2-way repeated measure ANOVA:<br>* DBS: $F_{14,336} = 1.764$, $P = 0.0427$.<br>* Group: $F_{1,24} = 26.01$, $P < 0.0001$.<br>* DBS x Group: $F_{14,336} = 0.9112$, $P = 0.5469$.<br><br>2-way repeated measure ANOVA on baseline sessions: |

| | | |
|---|---|---|
| | | * DBS: $F_{4,96} = 0.529$, $P = 0.7147$.<br>* Group: $F_{1,24} = 21.51$, $P = 0.0001$.<br>* DBS x Group: $F_{4,96} = 0.5776$, $P = 0.6796$.<br><br>2-way repeated measure ANOVA on ON sessions:<br>* DBS: $F_{4,96} = 3.367$, $P = 0.0127$.<br>* Group: $F_{1,24} = 15.29$, $P = 0.0007$.<br>* DBS x Group: $F_{4,96} = 0.654$, $P = 0.6255$.<br><br>2-way repeated measure ANOVA on OFF sessions:<br>* DBS: $F_{4,96} = 0.0848$, $P = 0.9869$.<br>* Group: $F_{1,24} = 21.05$, $P = 0.0001$.<br>* DBS x Group: $F_{4,96} = 1.653$, $P = 0.1673$.<br><br>Bonferroni *post-hoc* test:<br>* Sensitive vs. Resistant:<br>$P < 0.05$ for baseline 5 (power = 0.99).<br>OFF 4 (power = 0.96).<br>$P < 0.01$ for baseline 4 (power = 1).<br>ON 1,4,5 ($0.85 \leq$ power $\leq 0.95$).<br>$P < 0.001$ for ON 2 (power = 0.94).<br>OFF 2,3 ($0.98 \leq$ power $\leq 0.99$).<br><br>* Resistant:<br>$P < 0.05$ for ON 1 vs. Baseline 1-3,5 ($0.99 \leq$ power $\leq 1$).<br>ON 2 vs. Baseline 3 (power = 1).<br>$P < 0.01$ for ON 2 vs. Baseline 1,2,5 (power = 1). |
| **S2C** | Sensitive (n =14)<br><br>Resistant (n = 12) | 2-way repeated measure ANOVA:<br>* DBS: $F_{2,48} = 2.489$, $P = 0.0937$.<br>* Group: $F_{1,24} = 26.01$, $P < 0.0001$.<br>* DBS x Group: $F_{2,48} = 0.6766$, $P = 0.5131$.<br><br>Bonferroni *post-hoc* test:<br>* Sensitive vs. Resistant:<br>$P < 0.01$ for baseline (power = 0.99).<br>$P < 0.001$ for ON, OFF ($0.96 \leq$ power $\leq 0.99$). |
| **S2D** | | 2-way repeated measure ANOVA:<br>* DBS: $F_{14,336} = 1.718$, $P = 0.0505$.<br>* Group: $F_{1,24} = 15.84$, $P = 0.0006$.<br>* DBS x Group: $F_{14,336} = 0.9129$, $P = 0.545$.<br><br>2-way repeated measure ANOVA on baseline sessions:<br>* DBS: $F_{4,96} = 0.3151$, $P = 0.8672$.<br>* Group: $F_{1,24} = 11.64$, $P = 0.0023$.<br>* DBS x Group: $F_{4,96} = 0.2966$, $P = 0.8795$. |

| | | |
|---|---|---|
| | Sensitive (n =14)<br><br>Resistant (n = 12) | 2-way repeated measure ANOVA on ON sessions:<br>* DBS: $F_{4,96} = 2.042$, $P = 0.0946$.<br>* Group: $F_{1,24} = 8.251$, $P = 0.0084$.<br>* DBS x Group: $F_{4,96} = 1.1$, $P = 0.3612$.<br><br>2-way repeated measure ANOVA on OFF sessions:<br>* DBS: $F_{4,96} = 0.2549$, $P = 0.9060$.<br>* Group: $F_{1,24} = 12.14$, $P = 0.0019$.<br>* DBS x Group: $F_{4,96} = 0.9182$, $P = 0.4567$.<br><br>Bonferroni *post-hoc* test:<br>* Sensitive vs. Resistant:<br>$P < 0.05$ for OFF 4 (power = 0.83).<br>$P < 0.01$ for ON 2 (power = 0.83).<br>OFF 2 (power = 1).<br>* Resistant:<br>$P < 0.05$ for ON 2 vs. Baseline 4,5 ($0.99 \leq$ power $\leq 1$). |
| **S2E** | | 2-way repeated measure ANOVA:<br>* DBS: $F_{2,48} = 2.973$, $P = 0.0606$.<br>* Group: $F_{1,24} = 15.84$, $P = 0.0006$.<br>* DBS x Group: $F_{2,48} = 1.019$, $P = 0.3686$.<br><br>Bonferroni *post-hoc* test:<br>* Sensitive vs. Resistant:<br>$P < 0.01$ for ON, OFF ($0.8 \leq$ power $\leq 0.99$). |
| **S2F** | | Paired *t*-test:<br>*Sensitive: $t_{13} = 2.268$, $P = 0.041$ (power = 0.1).<br>*Resistant: $t_{11} = 3.576$, $P = 0.0043$ (power = 0.99). |
| **S2G** | | Paired *t*-test:<br>*Sensitive: $t_{13} = 2.115$, $P = 0.0544$.<br>*Resistant: $t_{11} = 2.414$, $P = 0.0344$ (power = 0.93). |
| **S3A** | Sensitive (n =14) | 1-way repeated measure ANOVA:<br>$F_{5,65} = 0.573$, $P = 0.7204$. |
| **S3B** | Resistant (n = 12) | 1-way repeated measure ANOVA:<br>$F_{5,55} = 2.099$, $P = 0.0793$. |
| **S4** | n = 6 | 1-way repeated measure ANOVA:<br>$F_{3,15} = 10.19$, $P = 0.0007$.<br><br>Bonferroni *post-hoc* test:<br>$P < 0.05$ for Saline-OFF vs. Cocaine-ON.<br>Saline-ON vs. Cocaine-ON.<br>$P < 0.01$ for Saline-OFF vs. Cocaine-OFF.<br>Saline-ON vs. Cocaine-OFF. |

| | | |
|---|---|---|
| **S5A** | 0 Hz (n =3)<br>30-Hz (n =3)<br>130-Hz (n =3) | 2-way repeated measure ANOVA:<br>* DBS: $F_{2,12} = 2.034$, $P = 0.1735$.<br>* Group: $F_{2,6} = 0.1722$, $P = 0.8458$.<br>* DBS x Group: $F_{4,12} = 0.1999$, $P = 0.9336$. |
| **S6A** | Sensitive (n = 36)<br><br>Resistant (n = 17) | Pearson correlation test:<br>$r = 0.1479$, $P = 0.2907$. |
| **S6B** | | 2-way repeated measure ANOVA:<br>* Session: $F_{4,204} = 0.9293$, $P = 0.4422$.<br>* Group: $F_{1,51} = 2.512$, $P = 0.1374$.<br>* Session x Group: $F_{4,204} = 0.5277$, $P = 0.7155$. |
| **S6C** | | 2-way repeated measure ANOVA:<br>* Session: $F_{4,204} = 1.823$, $P = 0.1258$.<br>* Group: $F_{1,51} = 2.278$, $P = 0.1191$.<br>* Session x Group: $F_{4,204} = 1.427$, $P = 0.2263$. |
| **S6D** | | 2-way repeated measure ANOVA:<br>* Session: $F_{4,204} = 3.587$, $P = 0.0075$.<br>* Group: $F_{1,51} = 2.844$, $P = 0.0979$.<br>* Session x Group: $F_{4,204} = 0.162$, $P = 0.9573$. |
| **S6E** | | 2-way repeated measure ANOVA:<br>* Session: $F_{4,204} = 10.71$, $P \leq 0.0001$.<br>* Group: $F_{1,51} = 1.241$, $P = 0.2705$.<br>* Session x Group: $F_{4,204} = 1.926$, $P = 0.1075$. |
| **S6F** | | Pearson correlation test:<br>$r = 0.05987$, $P = 0.6702$. |
| **S6G** | | Pearson correlation test:<br>$r = 0.09111$, $P = 0.5165$. |
| **S7C** | Sensitive (n =12)<br><br>Resistant (n = 5) | Pearson correlation test:<br>Theta band: $r = 0.6048$, $P = 0.0101$ (power = 0.83).<br>Beta band: $r = 0.5127$, $P = 0.0353$ (power = 0.63).<br>Gamma band: $r = 0.1869$, $P = 0.4726$. |
| **S7D** | | 2-way repeated measure ANOVA for sensitive:<br>* Frequency: $F_{62,693} = 0.5167$, $P \geq 0.999$.<br>* Sessions: $F_{1,693} = 12.21$, $P < 0.001$.<br>* Frequency x Sessions: $F_{62,693} = 1.091$, $P \geq 0.999$.<br><br>2-way repeated measure ANOVA for resistant:<br>* Frequency: $F_{62,252} = 1.548$, $P = 0.01$.<br>* Sessions: $F_{1,252} = 286.9$, $P < 0.001$.<br>* Frequency x Sessions: $F_{62,252} = 1.548$, $P = 0.01$.<br><br>Bonferroni *post-hoc* test:<br>$P < 0.05$ for 11, 17, 19 Hz.<br>$P < 0.01$ for 6, 10, 15, 16, 18 Hz.<br>$P < 0.001$ for 7, 8, 9 Hz. |

| | | |
|---|---|---|
| **S8A** | n = 5 STNs | 2-way repeated measure ANOVA:<br>* Frequency: $F_{138,556} = 6{,}131$, $P < 0.0001$.<br>* Sessions: $F_{2,1112} = 128.1$, $P < 0.0001$.<br>* Frequency x Sessions: $F_{276,1112} = 3.461$, $P < 0.0001$.<br><br>Bonferroni *post-hoc* test:<br>* Pre-8 Hz vs Post-8 Hz:<br>$P < 0.05$ for 16 Hz.<br>$P < 0.01$ for 15 Hz<br>$P < 0.001$ for 6-14 Hz.<br>* Pre-8 Hz vs Post-7 d:<br>$P < 0.05$ for 15 Hz.<br>$P < 0.01$ for 13-14 Hz<br>$P < 0.001$ for 6-12 Hz. |
| **S8B** | | 1-way repeated measure ANOVA<br>* Theta band: $F_{2,8} = 4.465$, $P = 0.0499$.<br>Pre-8 Hz vs. Post-8 Hz: $P < 0.05$ (power = 0.79)<br>* Beta band: $F_{2,8} = 2.846$, $P = 0.1165$.<br>* Gamma band: $F_{2,8} = 1.201$, $P = 0.1138$. |
| **S9A** | 0 Hz (n =5)<br><br>8-Hz (n =5)<br><br>70 Hz (n =3) | 2-way repeated measure ANOVA on escalation sessions:<br>* Sessions: $F_{14,140} = 3.136$, $P = 0.0003$.<br>* Group: $F_{2,10} = 0.3937$, $P = 0.6845$.<br>* Sessions x Group: $F_{28,140} = 0.9863$, $P = 0.4926$.<br><br>Bonferroni *post-hoc* test:<br>$P < 0.05$ for session 1 vs. sessions 13-15. |
| **S9B** | | 2-way repeated measure ANOVA:<br>* Sessions: $F_{13,140} = 0.601$, $P = 0.8502$.<br>* Group: $F_{2,140} = 1.079$, $P = 0.3426$.<br>* Sessions x Group: $F_{26,140} = 0.3683$, $P = 0.9979$. |
| **S9C** | | 2-way repeated measure ANOVA:<br>* Sessions: $F_{12,120} = 8.148$, $P < 0.0001$.<br>* Group: $F_{2,10} = 8.361$, $P = 0.0.0073$.<br>* Sessions x Group: $F_{24,120} = 0.7678$, $P < 0.0001$.<br><br>2-way repeated measure ANOVA on baseline sessions:<br>* Sessions: $F_{4,40} = 3.034$, $P = 0.0283$.<br>* Group: $F_{2,10} = 0.3723$, $P = 0.6983$.<br>* Sessions x Group: $F_{8,40} = 1.532$, $P = 0.177$.<br><br>2-way repeated measure ANOVA for punishment sessions:<br>* Sessions: $F_{7,70} = 9.131$, $P < 0.0001$.<br>* Group: $F_{2,10} = 9.532$, $P = 0.0048$.<br>* Sessions x Group: $F_{14,70} = 6.687$, $P < 0.0001$. |

| | | |
|---|---|---|
| | | Bonferroni *post-hoc* test:<br>* 0 Hz vs. 8 Hz:<br>$P < 0.05$ for session 6 (power = 0.68).<br>$P < 0.001$ for sessions 5,7-8 (0.83 ≤ power ≤ 0.98).<br><br>8 Hz:<br>$P < 0.05$ for session 4 vs. 8 (power = 1).<br>$P < 0.01$ for session 4 vs. 5 (power = 1).<br>$P < 0.001$ for session 1 vs. 5,7-8 (0.88 ≤ power ≤ 1).<br>session 2 vs. 5,7-8 (0.75 ≤ power ≤ 0.97).<br>session 3 vs. 5,7-8 (0.77 ≤ power ≤ 0.99).<br>session 4 vs. 7 (power = 1). |
| **S9D** | 0 Hz (n =5)<br><br>8-Hz (n =5)<br><br>70 Hz (n =3) | 2-way repeated measure ANOVA on baseline sessions:<br>* Sessions: $F_{2,10} = 1.791$, $P = 0.2164$.<br>* Group: $F_{1,10} = 3.276$, $P = 0.1004$.<br>* Sessions x Group: $F_{2,10} = 1.821$, $P = 0.2117$.<br><br>2-way repeated measure ANOVA on punishment sessions:<br>* Sessions: $F_{2,10} = 6.342$, $P = 0.0305$.<br>* Group: $F_{1,10} = 0.7365$, $P = 0.5030$.<br>* Sessions x Group: $F_{2,10} = 2.833$, $P = 0.106$.<br><br>Bonferroni *post-hoc* test:<br>* 8 Hz:<br>$P < 0.05$ for session 1 vs. 2 (power = 0.99). |
| **S9E** | | 2-way repeated measure ANOVA:<br>* Sessions: $F_{12,130} = 0.8742$, $P = 0.5747$.<br>* Group: $F_{2,130} = 0.9482$, $P = 0.3901$.<br>* Sessions x Group: $F_{24,130} = 0.8326$, $P = 0.69$. |
| **S10A** | n = 4 | 1-way repeated measure ANOVA:<br>$F_{2,6} = 3.34$, $P = 0.106$. |
